# Bacteria can anticipate the seasons: photoperiodism in cyanobacteria

**DOI:** 10.1101/2024.05.13.593996

**Authors:** Maria Luísa Jabbur, Carl Hirschie Johnson

**Author notes:** Each affiliation should be a separate paragraph. For large groups, use the name of the group or consortium and include a full list of the authors and affiliations at the end of the main manuscript or in the Supplementary Materials.

## Abstract

Photoperiodic Time Measurement is the ability of plants and animals to measure differences in day/night-length (photoperiod) and use that information to anticipate critical seasonal transformations such as annual temperature cycles. This timekeeping phenomenon triggers adaptive responses in higher organisms such as gonadal growth/regression, flowering, and hibernation. Unexpectedly, we discovered this capability in cyanobacteria, unicellular prokaryotes with generation times of only 5-6 h. Cyanobacteria in short winter-like days develop enhanced resistance to cold that involves desaturation of membrane lipids and differential programs of gene transcription, including stress response pathways. As in eukaryotes, this photoperiodic timekeeping requires an intact circadian clockwork and develops over multiple cycles. Therefore, photoperiodic timekeeping evolved in much simpler organisms than previously appreciated, and involved genetic responses to stresses that recur seasonally.

## Main Text

Many branches of the eukaryotic tree of life have evolved the ability to not only react to, but rather to anticipate, the changing seasons and proactively adjust their behavior and physiology by “measuring” seasonally changing photoperiods (PPs) with a timekeeping mechanism that often involves a circadian clock (*1*–*3*). However, prokaryotic organisms with very short generation times have barely been considered to harbor such a prolonged temporal program(*4, 5*). Cyanobacteria with doubling times as short as 5-6 h have nevertheless become productive model organisms for the study of circadian (daily) rhythms, with a core mechanism comprised of only three proteins that can be reconstituted *in vitro* (KaiABC(*6*–*9*)). Cyanobacteria are found in a wide range of latitudes, and as such are exposed to dramatic annual changes in light and temperature in their natural environment(*10*), and we therefore tested whether cyanobacteria are capable of timekeeping on a temporal base that is even longer than circadian, namely PPTM in promoting adaptive cold-resistance responses to seasonal changes.

### Short days promote cold resistance in cyanobacteria in a clock-dependent manner

Cells of the unicellular cyanobacterium *Synechococcus elongatus* PCC 7942 were grown at 30ºC on plates containing solid BG-11 medium and exposed to 8 continuous PP cycles of either short days (LD8:16, 8h of light followed by 16h of darkness), equinox (LD12:12) or long days (LD16:8). After this exposure, the plates were resuspended in liquid at the middle of their day phase and challenged with timed exposures to ice-cold temperatures, after which their survival was assessed (Fig. 1A). Prior exposure to short-days led to about 2-3X higher survival of wild-type cyanobacteria compared to those that were exposed to equinox or long days (Fig. 1B). This differential survival was not observed in arhythmic cells in which the circadian clock genes *kaiA, kaiB* and *kaiC* had been deleted (Δ*kaiABC*, Fig. 1B, fig. S2A and B), nor in mutants that lacked other components of the core clockwork or its output pathways (Δ*kaiA*, Δ*rpaA*, Δ*cikA*, Δ*sasA*, fig. S2A and C).

**Fig. 1.**
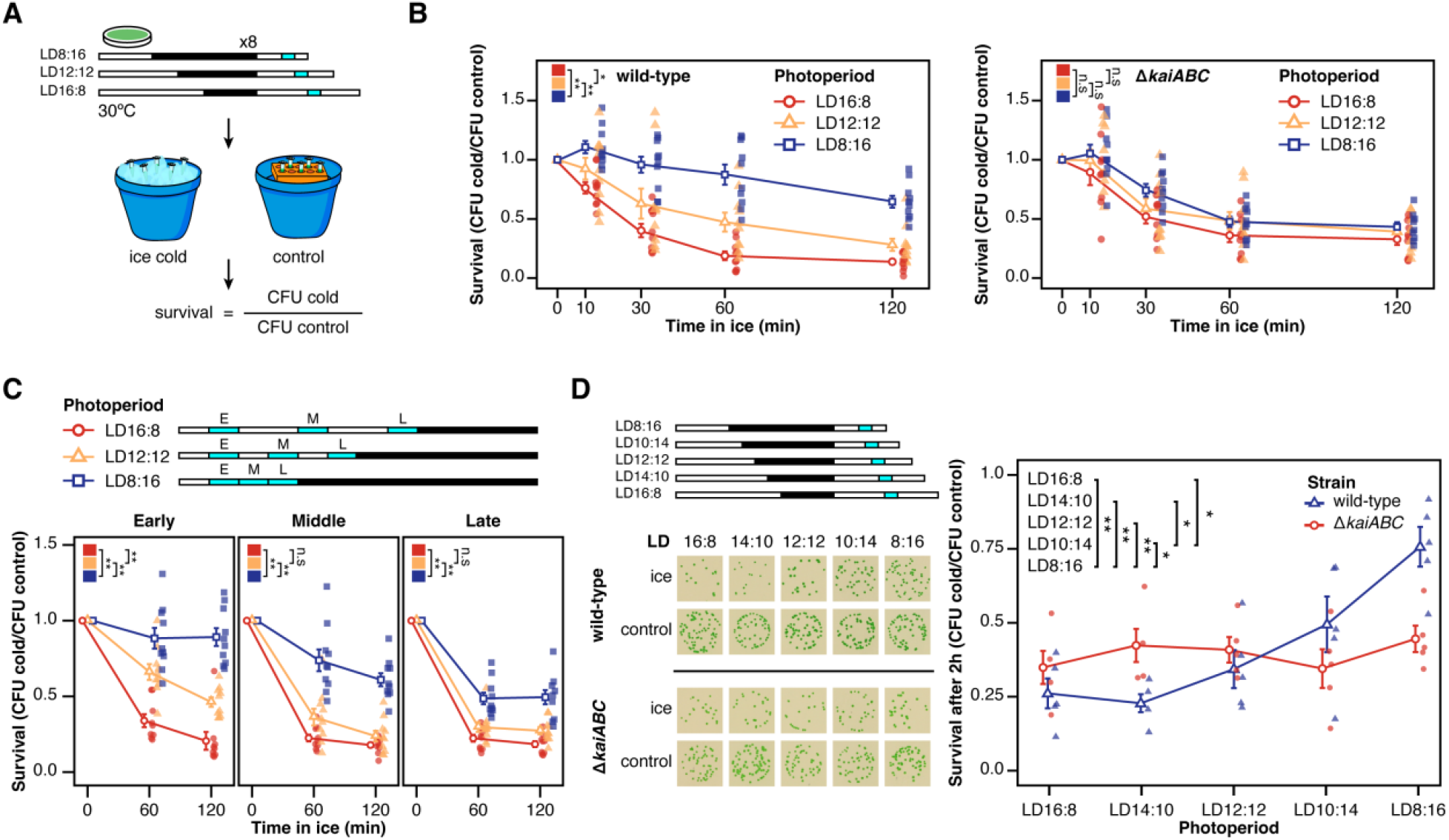
Cyanobacteria exposed to short days survive cold treatment better than those exposed to long days. (**A**) Experimental protocol used in the cold survival assay. See fig. S1A-E for details). Light blue bars indicate the phase (midday) in which the cold survival assay was performed. (**B**) Cold survival curve for wild-type (*n* = 10) and Δ*kaiABC* cells (*n* = 9 for LD16:8; *n* = 10 for LD12:12 except for 10min in which *n* = 9; *n* = 10 for LD8:16 except 120min in which *n* = 9). The results of statistical comparisons between photoperiods (determined by pairwise t-tests with Bonferroni correction) are expressed as asterisks (* < 0.05, ** < 0.01), with the comparison indicated by brackets around squares sharing a color scheme with the photoperiod data points (**C**) Similar to (B), but showing results for cells tested at early day (two hours after lights on, *n* = 10 for LD8:16 and LD12:12, *n* = 9 for LD16:8), midday (same as (B), *n = 10* except for LD16:8 at 60min, in which *n* = 9) and late day (two hours before lights off, *n* = 10, except for LD8:16 and LD16:8 120min, in which *n* = 9) (**D**) Original data on the left shows colony forming units for wild-type (top) and Δ*kaiABC* (bottom) cells exposed to a range of photoperiods as indicated, tested after 8 PP cycles at their respective middays. Right, quantification of the survival after 2h of exposure to ice (*n* = 5, 4, 5, 5, 5 for wild-type and *n* = 4, 4, 5, 5, 5 for Δ*kaiABC*). Significances levels refer to comparisons between wild-type values (there was no significant difference Δ*kaiABC*). For all graphs, error bars show the standard error of the mean, and open dots indicate the average.

To assess whether cold resistance was influenced by the time of day of the assay, we repeated the experiment in the morning, mid-day, and evening of the LD cycles (Fig. 1C for WT, fig. S3A for Δ*kaiABC*). The survival differential is highest when the assay is performed in the early day, but for all sampled times, WT survives cold best after exposure to short PPs as compared to long or equinox PPs; arhythmic Δ*kaiABC* cells show no PP or time-of-day differential survival (Fig. 1C, fig. S3B). We standardized our protocol to mid-day assays because that daily time is when the cells’ circadian phase should be most comparable across different PPs(*11*). In general, PPTM phenomena show differential responses along a continuum of PPs, and we therefore assayed along a gradient of PPs and found that long (LD16:8 and LD14:10) PPs lead to poor survival, while LD12:12 and shorter PPs progressively lead to increased cell survival (Fig. 1D). Survival of the Δ*kaiABC* cells is unaffected by photoperiod.

An intriguing aspect of PPic responses in general is that they are often history-dependent, i.e., an organism’s interpretation of a PP is dependent upon the prior exposure conditions(*2*). Consequently, a common characteristic of *bona fide* PPTM is that multiple cycles of inductive photoperiods are often required to elicit a complete response, namely there is a “photoperiodic counter”(*12*–*16*). Given their short generation time, can prokaryotic cyanobacteria remember and integrate a PP signal that extends over multiple cycles and generations? A single cycle of different PPs does not produce differential survival (Fig. 2A); at least 4 cycles seem to be necessary to achieve the full magnitude of the PPic response (Fig. 2B). Interestingly, the development of differential survival among PPs appears to be mainly caused by a progressive gain of resistance in short days, rather than by a further loss of cold resistance in long days. Therefore, the PPic response required cumulative exposure to multiple cycles, thereby involving a PPic counting “memory.” The short-day gain of cold resistance achieved by 4 PP cycles spanned at least ∼3.2 generations in WT cells (Fig. 2C). Again, cells without a functional circadian clock express a PP-independent increase in cold resistance as the number of LD cycles increases (fig. S4A and B). How long does this “memory” last after the PPic cues are removed? After our standard pre-treatment of 8 PP cycles, we transferred the cells to either constant light (LL) or constant darkness (DD) and measured thereafter at 24-h intervals. Transfer to LL leads to a general decrease in survival of the short-PP-exposed cells (Fig. 2D and fig. S4F), while the opposite is true for release to DD (fig. S4D and E). In both cases, after 24 h in constant conditions, short-PP-exposed cells tend to survive slightly more than cells from the other PPs, but these differences were not statistically significant (fig. S4F and G).

**Fig. 2.**
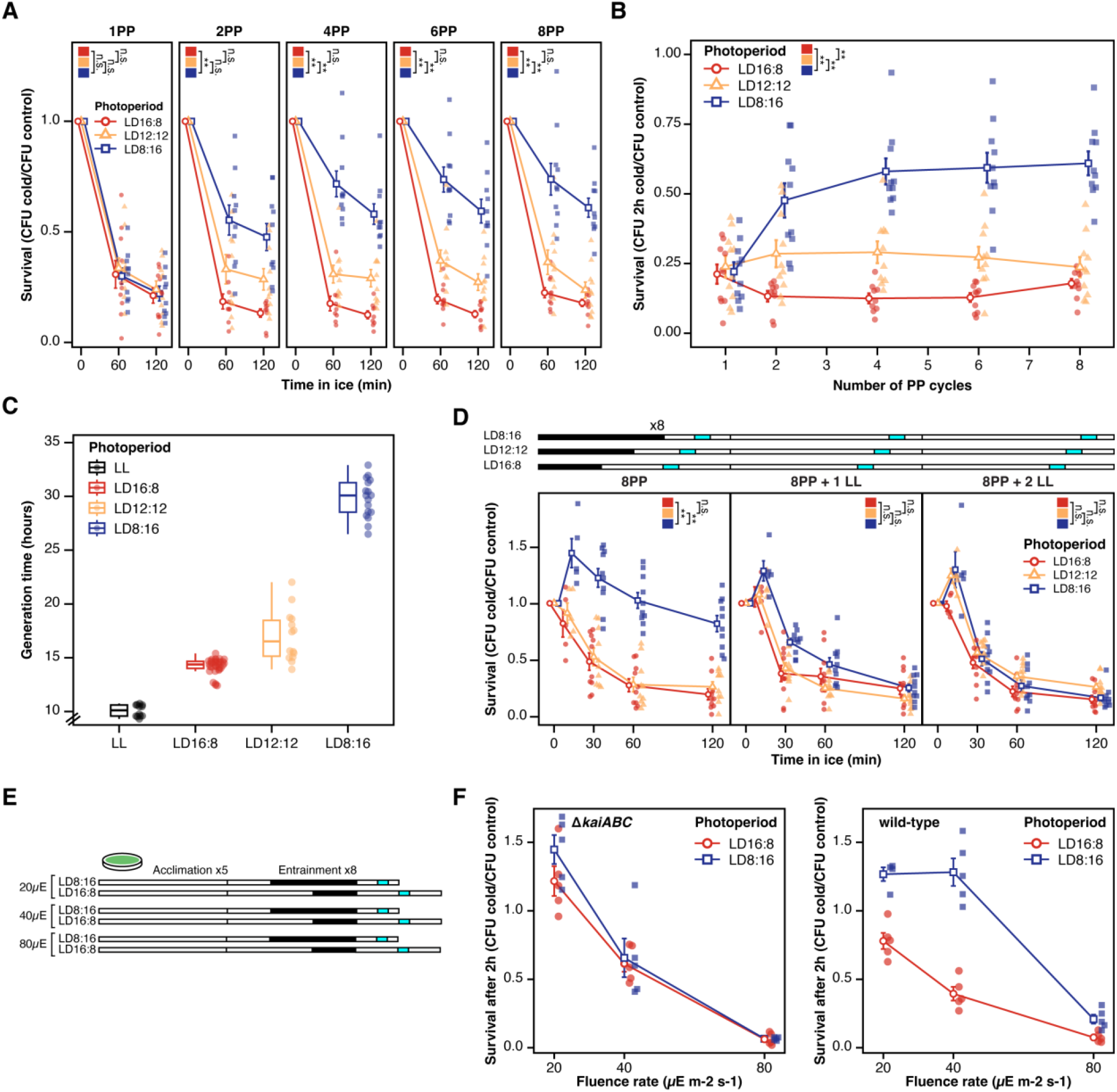
Cyanobacterial photoperiodic response takes multiple cycles to develop and spans multiple generations. (**A**) Survival curves similar to Figure 1b, but for cold survival assays performed after different numbers of PP cycles (*n* = 10 for all except LD16:8 at 2 PP and 4 PP, in which *n* = 9) (**B**) Survival after 2h of exposure to cold throughout the different number of PP cycles. (**C**) Generation time of cyanobacterial cells for each of the photoperiods analyzed, as well as LL (from left to right, *n* = 12, 32, 14 and 17). (**D**) Diagram and survival curves for the constant light assay (*n* = 10, except for 10min, in which *n* = 5). Blue bars indicate the times at which the survival assay was performed (midday, 24h and 48h after). (**E**) Diagram showing the protocol for the different light intensities and photoperiods tested. Blue bars indicate the time at which the survival assay was performed (**F**) Survival after 2h of exposure to cold at different fluence rates (*n=* 5 for all except for wild-type LD8:16 at 20µE, for which *n* = 4). Full survival curves, including other time points, can be found in fig. S5B and C. For all survival curves, open symbols denote the average and error bars denote the standard error of the mean. Significance was determined through pairwise t-tests with Bonferroni correction.

To be certain that our results were dependent upon the duration of the PP and not simply a differential exposure to light–which is critically relevant for a photosynthetic organism–we tested whether changes in light intensity could influence this photoperiodic response. Cells were exposed to fluence rates of 20µE m^-2^ s-^1^, 40µE m^-2^ s-^1^ (our standard light intensity), and to 80µE m^-2^ s-^1^ (Fig. 2E). Recovery of the Δ*kaiABC* cells from cold was a simple function of fluence rate: higher light intensities lead to lower survival than low light intensities (Fig. 2F; & fig. S5 for the full survival curves at each light intensity). On the other hand, while overall survival is also correlated with fluence rate for WT cells, in this case WT cells exposed to short days survived better than those exposed to long days at all intensities (Fig. 2F; fig. S5B), indicating that WT cells are integrating temporal information with light intensity. Taken together, these data indicate that the photoperiod-dependent response to cold exposure develops over multiple LD cycles that span multiple generations in cyanobacteria. Our data also suggest that cells in their natural environment integrate multiple factors (e.g., light intensity) along with photoperiod to program the optimal adaptive response.

### Short days lead to lipid membrane desaturation

Previous studies of cold acclimation in cyanobacteria have demonstrated that prior exposure to mild low temperatures can increase subsequent survival to colder temperatures, and have implicated lipid desaturation–thereby increasing membrane fluidity–as an important adaptation to cold(*17, 18*). In their natural environments, cyanobacteria experience a shortening of the daylength in concert with a lowering of temperatures as winter approaches. To confirm that photoperiod and prior temperature exposure are synergistic towards cold survival, we exposed cells to 8 cycles of either short, equinox or long days at constant 30ºC, followed by another cycle of each photoperiod at 20ºC, after which the cells were tested for their cold survival (Fig. 3A and B). Although cells exposed to 20ºC increased their survival in comparison to cells that did not receive the 20ºC treatment at all photoperiods, the cells exposed to short-days survived better than those exposed to the other photoperiods (Fig. 3B and C). Remarkably, in WT cells, the increase in survival caused by short-days without any 20ºC exposure was comparable to the effect of 20ºC exposure to the long-day cells (Fig. 3C). In “clockless” Δ*kaiABC* cells, 20ºC exposure enhanced cold survival, but different photoperiods did not (fig. S6A-C).

**Fig. 3.**
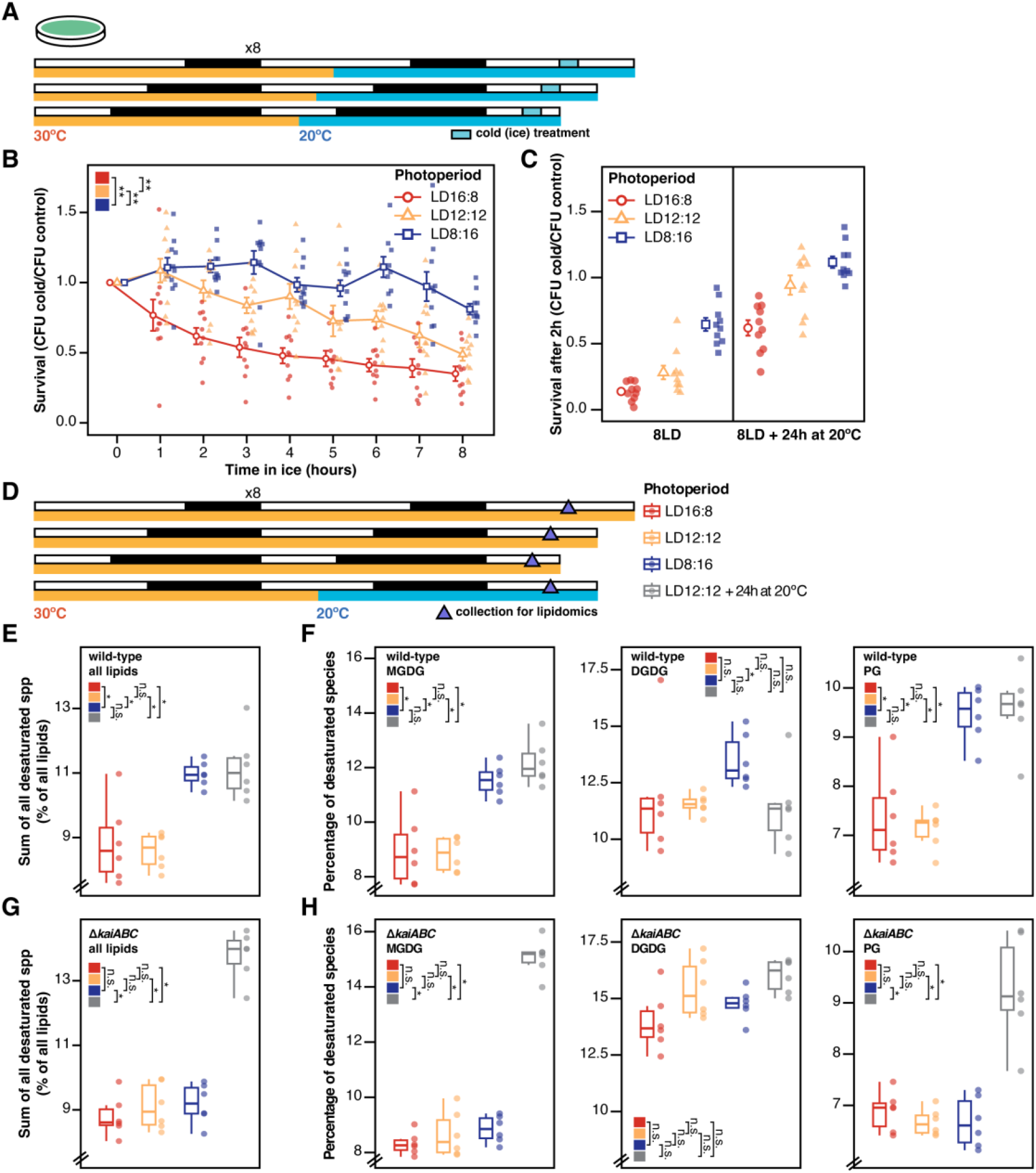
Exposure to short days leads to an increase in membrane desaturation akin to that resulting from exposure to low temperatures. (**A**) Diagram of the experiment testing the interaction of photoperiod and intermediate cold temperatures on survival. (**B**) Survival curve for wild-type cells treated as in a (*n* = 10 except for LD8:16 6h, in which *n* = 9). (**C**) Comparison between the survival of wild-type cells after 2h of exposure to cold, with and without prior treatment of 24h at 20ºC (*n* = 10). (**D**) Diagram of the lipidomics collection and prior treatment. (**E**) Sum of all desaturated lipid species for wild-type (*n* = 6, data in fig. S7, panels B {DGDG 30:2, 32:2 and 34:2}, C {MGDG 32:2 and 34:2} and E {PG 32:2 and 34:2}). (**F**) Percentage of desaturated species within each lipid class in wild-type (*n* = 6, data in fig. S8, panels A {DGDG 30:2, 32:2 and 34:2}, B {MGDG 32:2 and 34:2}, and C {PG, 32:2 and 34:2}). (**G**) Sum of all desaturated lipid species for Δ*kaiABC* (*n* = 6, data in fig. S9, panels B {DGDG 30:2, 32:2 and 34:2}, C {MGDG 32:2 and 34:2}, and E {PG 32:2 and 34:2}). (**H**) Percentage of desaturated species within each lipid class in Δ*kaiABC* (*n* = 6, data contained in fig. S10, panels A {DGDG 30:2, 32:2 and 34:2}, B {MGDG 32:2 and 34:2}, and C {PG, 32:2 and 34:2}). For panel (B), open symbols denote the mean and error bars denote the standard error of the mean. Significance was determined through pairwise t-tests with Bonferroni correction. MGDG = monogalactosyldiacylglycerol; DGDG = digalactosyldiacylglycerol; PG = phosphatidylglycerol.

Can short days in the absence of a temperature change mimic the molecular adaptive response of cold-induced lipid desaturation to anticipate winter? WT and Δ*kaiABC* cells were exposed to the 8 cycles of each photoperiod + 24h at 20ºC and extracted for lipidomic analyses (Fig. 3D). WT cells exposed to short days increased the percentage of desaturated lipid species in their cell membranes (esp. MGDG and PG, Fig. 3E and F). Notably, this increase was of the same magnitude as that observed for cells exposed to equinox + 20ºC for 24 h. Looking at individual lipid classes, we found that WT short-day and 20ºC-exposed cells share an increase in the desaturated species MGDG and PG, whereas short days increased the desaturated DGDG even more than 20ºC exposure (Figure 3F). In Δ*kaiABC* cells, however, only 20ºC exposure led to changes in lipid-saturation levels; different PPs in the absence of temperature changes had no effect in cells without the circadian clock genes (Fig. 3G and H). Apparently, short days elicit a *kaiABC-*dependent desaturation of membrane lipids in WT cells that could function to anticipate and adapt to seasonally cooling temperatures.

### Photoperiod determines global programs of gene expression

Do photoperiods differentially affect only lipid desaturation, or are other responses involved? To test whether distinct gene expression patterns underlie this photoperiodic adaptation, we performed RNA-seq on wild-type and Δ*kaiABC* cells that had been exposed to either 1 PP cycle (sufficient to synchronize circadian oscillations(*19*) but not to evoke the photoperiodic response; Fig. 2A and B), 4 PP cycles (photoperiodic response has developed; Fig. 2A and B), and 8 PP cycles (our usual testing condition)(Fig. 4A). Unexpectedly, we found that global programs of gene expression are differentially elaborated between short- and long-day exposed cells (722 genes = ∼26% of the genome at 8 PP), and over half of the differentially expressed genes achieve that status by 4 PP (Venn diagrams in Fig. 4B, volcano plots in Fig. 4C, PCA & distance matrix in fig. S11). Comparing the expression of WT short-day exposed cells to that of WT long-day cells in each of the PP cyclic conditions, there is no obvious correlation between the log_2_(fold change) at 1 PP cycle versus 8 PP cycles (R^2^_adj_ = < 0.01; Fig. 4B, left panel), but the differences in expression at 4 PP are well-correlated with those at 8 PP (R^2^_adj_ = 0.44, p < 0.001; Fig. 4B, right panel). This means that the changes in gene expression at 1 PP are likely to reflect immediate differences in entrainment, while 4-8 PP cycles are necessary to fully develop the PP-specific genetic programs. Gene expression of Δ*kaiABC* cells in short vs. long days is significantly different, but the number of differentially affected genes is almost 2X greater in WT cells (722 in WT, 401 in Δ*kaiABC* at 8 PP, and the overlap between these groups is only 288 genes, fig. S12). Therefore, many more genes are differentially expressed between short days and long days in WT but not in Δ*kaiABC* cells.

**Fig. 4.**
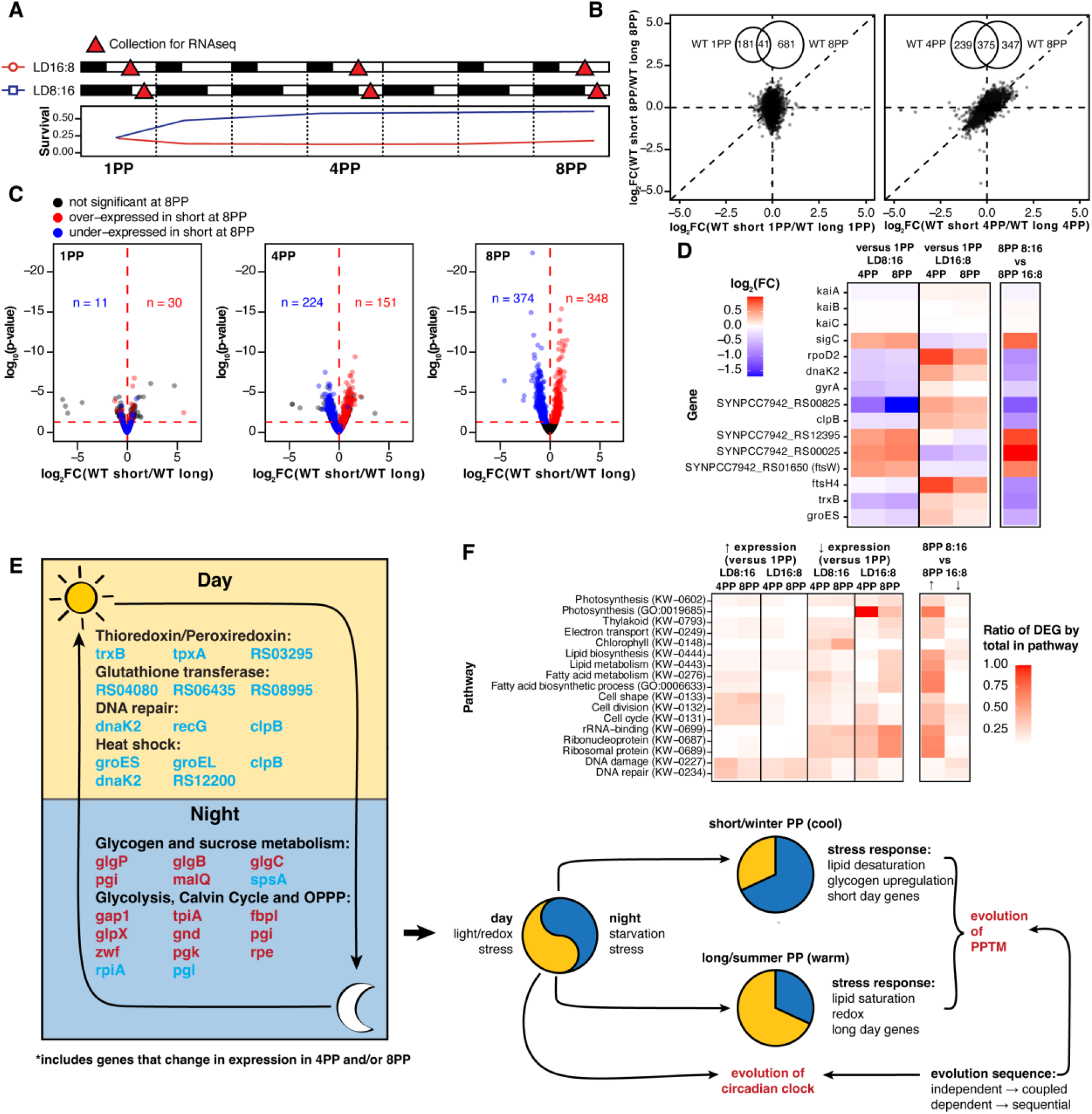
Exposure to short and long days leads to global alterations in gene expression that are evident after multiple PP cycles. (**A**) Diagram showing timepoints collected for RNA-seq (*n* = 3). (**B**) Correlation plot showing the log_2_(fold change) in expression of short-day cells divided by long-day cells, comparing either 8 PP to 1 PP (left) or 8 PP to 4 PP (right). The Venn diagrams show the number of differentially expressed genes shared between each condition (not including genes differentially expressed in opposite ways). Vertical and horizontal dashed lines indicate the y and x zero intercepts, and the diagonal line shows y=x. (**C**) Volcano plots showing the differentially expressed genes for wild-type at 1 PP, 4 PP and 8 PP, colored by the difference in expression at 8 PP. The numbers in blue on the bottom plot refer to the total number of genes under-expressed in short-days for wild-type after 8 PP, while those in red show the genes over-expressed. On the middle and top plots, the numbers refer to genes under or over-expressed in wild-type after either 4 PP or 1 PP that are also under or over-expressed in wild-type after 8 PP. (**D**) Subset of genes of interest (fig. S13), showing the circadian clock genes *kaiA, kaiB* and *kaiC* (which did not show significant changes in expression) as well as genes related to DNA metabolism, fatty acid metabolism and stress. The first four columns show the log_2_(fold-change) in expression in wild-type cells within each photoperiod of either 4 PP or 8 PP against 1 PP. The remaining column on the right shows the resulting differential expression between wild-type cells exposed to 8 PP of short-days vs 8 PP of long-days. (**E**) Model of the evolution of adaptive PPTM in relationship to the circadian clock. Stresses that occur primarily in the day vs. the night over the daily cycle are expanded and/or contracted as the daily photoperiod changes. This leads to differential stress response activation over the annual cycle whence a PPTM mechanism could evolve. (**F**) Similarly to (D), this panel shows a selection of the pathways of interest identified by gene ontology analysis within our dataset of genes of interest. Here instead of log_2_(fold-change), we show the ratio of differentially expressed genes (DEG) by the total amount of genes that annotated as part of that pathway. The first 4 columns show the ratio of over-expressed genes within each pathway, and are akin to the first four columns in panel (D). The 4 columns afterwards show the ratio of under-expressed genes. The remaining two columns on the left show the resulting ratio of over and under-expressed genes. KW refers to UniProt Keywords, and GO refers to Gene Ontology annotation.

Some genes of particular interest are highlighted in Fig. 4D, especially genes related to transcription, fatty acid metabolism, cell cycle, and stress responses. In prokaryotes, sigma factors regulate many genes synergistically to respond to environmental conditions(*20*) and we observed that the sigma factors *rpoD2* and *sigC* were differentially triggered; long days stimulated *rpoD2* expression, while short days promoted *sigC* (Fig. 4D). Both *rpoD2* and *sigC* are direct targets of the circadian transcriptional factor *rpaA*(*21*) and could relay circadian and photoperiodic information to the rest of the genome. We tested whether sigma factors were directly related to the cold-sensitivity response by performing the photoperiodic assay in knock-out cells of the sigma factors, *rpoD2, rpoD3, rpoD4*, and *sigC* (fig. S2D); while there were no significant differences in the cold-sensitivity response of Δ*sigC*, Δ*rpoD3*, or Δ*rpoD4*, all cells of the Δ*rpoD2* strain regardless of photoperiodic treatment became cold sensitive, suggesting that *rpoD2* might play a role in PP output such that deleting *rpoD2* short-circuits the photoperiodic response. Many of the genes of interest from our RNA-seq analyses have been implicated in stresses that occur in the day (bright light, UV, redox stress, hotter temperatures) versus night (metabolic stress) over the daily cycle (Fig. 4E).

Gene ontology analysis through DAVID(*22, 23*) identified differentially controlled pathways of interest, including the key processes of lipid/fatty acid metabolism, photosynthesis, cell cycle, DNA recombination & repair, and ribosomal activity (Fig. 4F). Overall, the RNA-seq results show that exposure to short vs. long days led to distinct programs of gene expression, caused partially by entrainment itself (changes common to cells exposed to 1 PP and 8 PP), and partially by direct responses to short days/long nights (changes common to WT and Δ*kaiABC* cells). However, there are at least 717 genes that are regulated at the behest of the *kaiABC*-dependent PP timekeeping mechanism (i.e., those that are WT specific for exposure to either 4/8 PP cycles; Tables S1 and 2 and fig. S13). This group of PPTM-regulated genes include those related to (i) DNA metabolism (*dnaK, gyrA, clpB*, the sigma factors *sigC* and *rpoD2*, and the class I SAM-dependent methyltransferase SYNPCC7942_RS00825), (ii) temperature (the chaperone *clpB* is a heat-shock protein related to thermotolerance in cyanobacteria(*24*), but it is also strongly induced by cold(*25*)), (iii) photosynthesis (thylakoid, electron transport, chlorophyll, photosynthesis(*26*)), and (iv) stress responses. In the stress response category, stimulation by long days include multiple genes/pathways that are associated with the light/heat stress that accompanies longer exposure to light, namely ftsH4 (high light(*27*)), trxB (oxidative stress(*28*)), groES (heat shock(*29*)), and the “stringent response” of cyanobacteria (redox stress mediated through changes in (p)ppGpp(*30, 31*) fig. S14). Survival in darkness is associated with glycogen in cyanobacteria(*32*), and we observed that the gene encoding the rate-limiting step for glycogen synthesis (glgC, encoding glucose-1-phosphate adenylyltransferase) is over-expressed in short days (fig. S15). Similarly, we see that multiple gene-related pathways associated with glycogen metabolism (glycolysis, sucrose metabolism, the Calvin cycle and the oxidative pentose phosphate pathways) were differentially expressed between short and long days (fig. S15), indicating that exposure to different photoperiods regulate genes encoding crucial metabolic enzymes. Interesting, the clock genes *kaiA, kaiB, kaiC* and *rpaA* were not differentially expressed between short and long-days at any of the PP cycles measured, implying that stability of the core circadian clockwork is conserved while photoperiod is changing seasonally. Finally, expression levels prominently changed for genes related to lipid & fatty-acid metabolism and membrane transport, such as the fatty acid desaturase SYNPCC7942_RS12395, the permease SYNPCC7942_RS00025, and the flipase FtsW (SYNPCC7942_RS01650(*33, 34*)). These genetic changes are consistent with our observation of molecular increases in the desaturation of membrane lipids, particularly in MGDG and PG levels (Fig. 3E-H, figs. S6-10).

## Discussion

These results show that cyanobacteria exposed to winter-like photoperiods are capable of surviving low temperatures 2-3X better than those exposed to summer-like photoperiods, and that this response requires a functional circadian clock. Short vs. long days promote distinct transcriptional programs, and cells exposed to short days experience an adaptive change in the saturation of membrane lipids akin to that seen in cells exposed to cool temperatures. The amplitude of the cold photoperiodic survival can be modulated by several environmental factors– namely, the daily phase of testing, light intensity, and previous exposure to cooler temperatures. In nature, as winter approaches, organisms face the shortening of daylength (and lower light intensity**(*35*)**) that precedes a gradual lowering of average temperature by ∼1-2 months**(*36, 37*)**). In their natural environment, cyanobacteria likely integrate multiple environmental factors to establish the timing and magnitude of their photoperiodic response. At first, it seems surprising that an organism with such a short generation time has evolved a mechanism to keep track of seasons and adaptively respond pre-emptively, but when selection is considered to be acting upon the population lineage rather than on the individual, the advantage of PPTM to cyanobacteria becomes intelligible**(*38, 39*)**.

These observations indicate that PPTM may have more ancient evolutionary roots than previously appreciated; it may have evolved before multicellularity and before the evolution of eukaryotes. We were particularly struck by the observation that short vs. long photoperiods induced counterbalancing responses in stress pathways whereby short days induced genes that confer adaptive responses to cold temperatures (e.g., lipid desaturases and glycogen synthesis), while long days induced stress response pathways associated with light/redox/heat stress (e.g., the “stringent response,” trxB, etc.). Of interest, when our RNA-seq dataset is compared with an independent dataset produced by exposing *S. elongatus* PCC 7942 to salinity & cold stressors for 1 h and 24 h**(*40*)**, the genes that were up-regulated in the other study after 24 h of increased salinity or lower temperature (20ºC) are also up-regulated in our dataset by short days after 8 PP, and a similar correlation is seen for down-regulated genes (ditto for Δ*kaiABC*, although at smaller fold-changes, fig. S18). These associations lead us to propose that PPTM might have first evolved in prokaryotes from pre-existing stress pathways that had originally evolved to combat acute stresses.

For obligate photoautotropic organisms like cyanobacteria, stresses that occur primarily in the day (bright light, UV, redox stress, hotter temperatures(*31*)) vs. the night (metabolic stress, starvation) over the daily cycle are expanded and/or contracted as the daily photoperiod changes (Fig. 4E). This leads to differential stress pathway activation over the annual cycle. Once a stress pathway exists, the introduction of a timekeeping mechanism to additionally regulate the stress pathway to not only respond acutely but also to anticipate regularly occurring environmental stresses is a logical selective pressure to evolve a PPTM. While photoperiod has been previously labeled as a possible stress in photosynthetic eukaryotes(*41*), the hypothesis that PPTM may have evolved from pre-existing stress response pathways has not been advanced prior to this study. Moreover, our results raise the question of which evolved first: a circadian or a PPTM system? While PPTM is now often associated with a circadian timekeeper, the first photoperiodic timing mechanism might have been an “hourglass timer” that later became linked to a pre-existing circadian timer OR the hourglass PPTM timer might have itself been the ancestor of a more flexible self-sustained circadian system (Fig. 4E). Therefore, the evolutionary sequencing between the daily circadian clock and the seasonal PPTM might be two independent timekeepers that became coupled, or the sequential evolution of one from the other (Fig. 4E).

The demonstration of PPTM in cyanobacteria has economic and global implications. Summer “blooms” of cyanobacteria in lakes and ocean can be economically devasting. These blooms have usually been interpreted to result from acute, local conditions (temperature, light intensity, nutrients), but perhaps they are also a manifestation of an anticipatory seasonal/PPic response. Finally, for all organisms that respond adaptively to photoperiodic changes–now including cyanobacteria–the spectre of climate change/global warming means that the relationship between average temperature and any particular photoperiod is rapidly shifting; cyanobacteria may be able to evolve rapidly enough to alter their anticipatory PPic responses, but organisms with longer generation times that cannot change their latitudinal range may be trapped by seasonal responses that are no longer timed appropriately**(*42*)**. Altogether, these results provide the first report of *bona fide* adaptive photoperiodic timing in a prokaryote, opening the possibility that photoperiodism might be an evolutionarily ancient phenomenon, as well as introducing a new model system to study the mechanisms and evolution of photoperiodic responses.

## Acknowledgments

We thank Dr. Yao Xu, who generated many of the strains used in this manuscript, as well as Dr. Hideo Iwasaki, who gave us the wild-type and Δ*kaiABC* strains used in most of the experiments reported. We also thank the multiple chronobiologists who have through the years offered input on this project. We are grateful to Angela Jones of Vanderbilt Technologies for Advanced Genomics (VANTAGE), who helped us with their professional sequencing services, and Ruth Welti and the Kansas Lipidomics Research Center facility who aided us with lipidomics.

## Funding

National Institutes of Health grants GM107434 and GM067152 (CHJ)

Vanderbilt University’s College of Arts and Science Summer Research Award (MLJ)

Vanderbilt University’s Russell G. Hamilton Graduate Leadership Institute Dissertation Enhancement Grant (MLJ)

National Institutes of Health grant P20GM103418 (Lipidomics Research Center)

National Science Foundation grant DBI-1726527 (Lipidomics Research Center)

United Stated Department of Agriculture Hatch/Multi-State project 1013013 (Lipidomics Research Center)

## Author contributions

Conceptualization: MLJ, CHJ

Methodology: MLJ, CHJ

Investigation: MLJ

Visualization: MLJ

Funding acquisition: CHJ

Project administration: CHJ

Supervision: CHJ

Writing – original draft: MLJ and CHJ

Writing – review & editing: MLJ and CHJ

## Data and materials availability

RNA-sequencing data have been deposited in the Gene Expression Omnibus under accession GSE252562. The processed data and statistics for all transcripts is available in Tables S3 and 4. The annotations obtained from DAVID are available in Table S5.

