## Supplemental figures for "Bacteria can anticipate the seasons: photoperiodism in cyanobacteria"

### Materials and Methods

#### Strains

Throughout the paper, the strains "wild-type" (WT) or "Δ*kaiABC*" refer to strains that harbor a bacterial luciferase P*kaiBC::luxAB* reporter targeted to Neutral Site I (NSI) to allow the monitoring of circadian gene expression (as in fig. S2B). These strains were a gift from Dr. Hideo Iwasaki(*43*, *44*). The wild-type strain harbors a chloramphenicol resistance (Cm) cassette, while the Δ*kaiABC* strain harbors both chloramphenicol (Cm) and kanamycin (Kn) resistance. We have separately tested strains with different resistances to ensure that antibiotic resistance did not play a role in the photoperiodic response described here (fig. S1F). The Δ*sigC*, Δ*rpoD2*, Δ*rpoD3* and Δ*rpoD4* all harbored spectinomycin resistance (Sp) and had a P*psbAI::luxAB* reporter on NSI. Inactivation of each gene was achieved by inserting either a gentamycin (Δ*sigC* and Δ*rpoD4*), a kanamycin (Δ*rpoD2*) or a chloramphenicol resistance marker (Δ*rpoD3*).

#### Measurement of luminescence rhythms

Cells were prepared and luminescence rhythms measured as described elsewhere(*45*). Luminescence intensity reports the activity of the *kaiBC* promoter (P*kaiBC::luxAB* reporter(*6*)). Prior to release into constant light for the luminescence assay, plates containing the wild-type and Δ*kaiABC* strains were given 12-h dark pulses for 3 consecutive days to synchronize the rhythms of the cells in the population. WT cells express robust circadian rhythms of luminescence, whereas Δ*kaiABC* cells are arhythmic (fig. S2B).

#### Culture Conditions

Bacterial cultures were streaked and grown in plates containing modified solid BG-11 medium(*46*) with 1 mM sodium thiosulfate(*47*). No antibiotics were added to solid or liquid media during experiments.
Unless otherwise noted, all experiments were performed with cyanobacteria cultures that were incubated in rooms kept at 30ºC under cool-white fluorescent light (~40µE $m^{-2}$ $s^{-1}$, see fig. S16A for the spectrum of the lights used). To standardize the environmental conditions among the photoperiod treatments (temperature, etc.), all photoperiodic treatments were conducted simultaneously and within the same room. Therefore, the temperature-controlled room was maintained on LD16:8, and shorter photoperiods were created by placing the plates at the end of their "day" phase in a light-tight box kept in the same room. We observed no relevant differences between the temperatures inside or outside of the light-tight box during the dark phase; the temperature difference in the room between lights-on and lights-off was less than 1.5ºC (fig. S16B). Based on our experience conducting these experiments in different temperature-control rooms, we advise experimenters who wish to repeat these experiments that they carefully monitor temperature and/or perform their own cold survival curve and choose the duration of cold treatment that best suits their experimental setup, as we have found that small changes in

average temperature can impact overall survival and potentially how many cycles it takes for the response to fully form (fig S17) (*48*).

Photoperiodic time-measurement experiments

Plates with cyanobacteria were kept in environmental conditions as described in the previous section. In order to account for the fact that plates kept in LD16:8 generally grow faster than those in LD12:12 or LD8:16 (due to the different durations of light to which the photoautotrophic cells were exposed), we decided to normalize their growth by challenging the plates with cold after they had been in 60h of light total. Thus, the cells exposed to LD16:8 and LD12:12 (or any other photoperiodic cycle with a day longer than LD8:16) were re-inoculated into fresh plates in a manner that allowed them to be exposed to approximately 60h of light prior to the cold challenge. This allowed us to have cells that had seen the same number of LD cycles but that were generally matched in their growth stage (fig. S1A). Since this introduced the possibility that the levels of desiccation of the plates might vary among the different photoperiods, we performed controls in which we re-inoculated LD12:12 and LD16:8 cells both on freshly-made plates and on plates that had been left in the experimental room since the onset of the experiment, and we found this comparison had no impact on the results (fig. S1B).

In the case of experiments in which we tested cells that were exposed to less than 8 photoperiodic cycles (8 "PP cycles," Fig. 2A and B, fig. S4A and B), light was supplemented prior to the beginning of the experiment to a total of 60h by putting the plates in a room that was kept in constant light (LL) at 40 µE $m^{-2}$ $s^{-1}$ and constant 30ºC. For experiments that tested different light intensities (Fig. 2F, fig. S5), the cells received an additional acclimation of 5 days prior to the photoperiodic entrainment, in which they were exposed to constant light at either 20, 40 or 80µE $m^{-2}$ $s^{-1}$ (fig. S18A). These light intensities were tested simultaneously, and as such the different light intensities were achieved by using combinations of light filters (LEE Filters: 250 (Half White Diffusion), 450 (Three Eight White Diffusion), 416 (Three Quarter White Diffusion) and 216 (White Diffusion). The spectra of the combinations of fluorescent + LED lights, as well as that of each filter, were practically identical, as shown in fig. S18D.

After treatment with photoperiods (PPs) for the specified number of photoperiodic cycles (unless otherwise specified, that number was 8 PP cycles), the cells on each plate were resuspended in ~1.5ml of liquid BG-11 medium at 30ºC. The $OD_{750}$ of each cell suspension was measured using a Shimadzu UV-1800 spectrophotometer and the suspensions were then diluted to an $OD_{750}$ of $10^{-4}$ (~$2.75x10^4$ CFU/ml). This dilution was then divided into two microcentrifuge tubes, one of which was put onto ice (0ºC) and the other which was maintained at 30ºC (the control). After different intervals of time, 2 µL of each of the experimental and control samples were spot-plated on BG-11 solid medium. This entire procedure was performed in a laminar-flow hood kept inside of an environmental control room maintained at 30ºC. In the case of the constant darkness (DD) experiments, the cells were processed under a dim green safe-light. All plates were then grown for 5 d in LL at 30ºC, after which time, the plates were photographed and the number of colony-forming units (CFUs) from each spot was determined. From these data, we derived a value for “survival,” which was the ratio of CFUs observed in the experimental samples that had been challenged with cold versus the CFUs of their respective control plates.

In some of the figures (particularly at timepoints in which cells were exposed to short durations of cold), the "survival" value was above 1. This is due to an inherent issue that emerges when pipetting liquids that are much colder than the ambient temperature(*49*), specifically that colder liquids tend to be over-pipetted. As this factor allowed for better visualization of the differences between photoperiods, we presented the data in the main figures as un-corrected for this phenomenon. However, we have measured conversion factors by weighting the volume of BG-11 media pipetted with a pipette set to 2 µL over the different durations of ice exposure (fig. S1C), and survival curves that are corrected for this factor are depicted in fig. S1D.

Growth curves

To measure the growth of cyanobacteria in solid media, we adapted a previous method(*50*). Namely, a lawn of cyanobacteria was first grown for 2 d in LL, after which the cells were resuspended and their optic density ($OD_{750}$) was measured. From these primary plates, we inoculated secondary plates with 100µL of the resuspended cyanobacterial cells in BG-11 medium (cell density was $OD_{750}$ = 3), which were then treated as described above for the experiments with different light intensities. Separately, we inoculated a set of six plates with the cyanobacterial solution, and 5 min after inoculation the plates were resuspended to allow us to estimate what proportion of the cells we recover with our collection/resuspension methodology (fig. S18A). This procedure allowed us to normalize our starting inoculation values. During LL and LD, the plates were resuspended twice with 750 µL of BG-11 after 30 ~ 80 h of cumulative light exposure (shorter intervals for plates exposed to high light intensity; longer intervals for low intensity). The optic density of the suspension was measured using a Shimadzu UV-1800 spectrophotometer, and the number of generations that had occurred since initial inoculation was established as the $\log_2$ of the ratio between the measured OD and the initial normalized starting inoculum. Generation time was calculated by dividing the number of generations that had elapsed by the total amount of hours (both light and dark) since inoculation.

As this assay required that we inoculate plates with a liquid suspension—unlike our initial assay that involved streaking bacteria in plates—we sought to evaluate whether this procedure affected our results and establish what optic density to use. We tested initial inoculums of 100µL of $OD_{750}$ = 2 ($5.5x10^8$ CFU/ml) and of $OD_{750}$ = 3 ($8.25x10^8$ CFU/ml), both of which yielded the same photoperiodic differences we saw with the streaked plates. As the "$OD_{750}$=3" plates had an overall higher survival than the "$OD_{750}$=2" plates (fig. S18C), we performed the rest of the growth curves and associated light intensity assays with "$OD_{750}$=3" inoculums.

Lipidomics

Approximately 1.4 ml of a cell suspension at an $O.D._{750}$ ~10 was centrifuged for ~1 min, after which we removed the resulting supernatant and measured the weight of each sample, which was then stored at -80ºC until used. Lipids were extracted from the cell pellet with the Blight and Dyer method of extraction(*51*). After extraction was complete, samples were maintained in a solution of 2:1 chloroform:methanol with 0.05% butylated hydroxytoluene (BHT) at -20ºC. The samples were shipped on dry ice to the Kansas Lipidomics Research Center Analytical Laboratory, where they were processed with a Xevo TQ-S triple-quadrupole mass spectrometer (Waters Corporation, Milford, MA, USA) and the total as well as relative amount of each lipid

species was determined. Our analyses only included samples whose coefficient of variation (standard deviation/average) made by pooling samples was less than 0.3.
Data were expressed as either the percentage that a given lipid species (e.g. MGDG 32:2) represented among all lipids quantified, or as the percentage of desaturated lipids within a lipid class (e.g. MGDG). These percentages were calculated by summing all of the more desaturated lipid species within each class or across all classes (e.g. we would sum the percentage of MGDG 30:2 and MGDG 32:2, but would not consider the percentage of MGDG 30:1 and 32:1, as these represent the less desaturated forms). When shown within class, the sum was weighted by the total amount of that particular class, and as such varies between 0 and 100% regardless of how abundant a lipid class was in any given sample.

RNA-seq

Samples of approximately 1.4 ml of a cell suspension at an $O.D._{750}$ ~10 (collected from a single bacterial plate) were centrifuged for 5 min at 10,000 rpm, after which the resulting supernatant was removed and the pellet samples were flash-frozen in liquid nitrogen and stored at -80ºC for future analyses. All cells were harvested at the middle of the day of their respective photoperiods. Cell pellets were later resuspended with BG-11 and mixed with RNAprotect Bacteria Reagent (Qiagen). Total RNA extraction was achieved using the RNeasy Mini Kit (Qiagen), and DNA contamination was removed with DNase I (RNase-free) (New England BioLabs). The samples were then submitted to Vanderbilt University Medical Center's VANTAGE Core, who determined the samples' integrity (RIN), depleted the rRNA (NEBNext rRNA Depletion Kit {Bacteria}), prepared the cDNA library, and sequenced the samples. All samples used had a RIN $\geq$ 8.1 and yielded at least 27M reads. Sequencing was performed on an Illumina NovaSeq 6000, with paired-end 150 bp.
Sequence reads were pre-processed to remove adaptors, low quality reads and reads shorter than 20 bp using TrimGalore v. 0.6.10(*52*). Indexes were built with hisat2 v. 2.2.1(*53*) using GenBank assembly accession GCA_000012525.1. hisat2 was also used to align our sequencing reads with the genome, and samtools(*54*) v. 1.11.0 was used to convert the files to .bam. These were mapped to the genome annotation file obtained from NCBI, and StringTie v. 2.1.7(*53*) was used to estimate transcript abundance. Finally, the prepDE.py script provided with StringTie was used to generate a matrix of raw read counts, which was then analyzed with DESeq2 v.1.36.0(*55*). The heatmap (fig. S11A) showing the calculated Euclidean distance between each of the samples ("stats" package(*56*)) was created with the package pheatmap(*57*) v. 1.0.12, normalized using the RLD (relative local distance) method. P-values were calculated by DESeq2 using a Wald test adjusted for multiple comparisons. To account for high levels of variation due to low counts or high dispersion, the $\log_2$(fold-change) values were shrunk with adaptive shrinking ("ashr" package(*58*) v.2.2.54).
To identify genes of interest, we created two subsets: namely, (i) one subset with genes that were differentially expressed between short-day and long-day WT cells but were either not differentially expressed or were differentially expressed in the opposite direction between Δ*kaiABC* short-day and long-day cells at 8 PP and WT short-day and long-day cells at 1 PP (fig. S13A, Table S1) , and (ii) another subset obtained by comparing the expression after 8 PP versus 1 PP cycle of the same photoperiod, and selecting genes that were differentially expressed (but not in the same direction) after 8 cycles of either short or long day PPs in wild-type, but whose

expression did not change (or changed in an opposite way from WT) between 1 PP and 8 PP for Δ*kaiABC* (see fig. S13B, Table S2). The genes highlighted in Figure 4d were taken from the combination of the two subsets described.
Gene ontology was performed with DAVID(*22*, *23*) (Database for Annotation, Visualization and Integrated Discovery). All current annotations available for the genes of *Synechococcus elongatus* PCC 7942 were extracted from DAVID, and genes deemed of interest (fig. S13, Tables S1 and 2) were given as input to identify potential pathways that are up- or down-regulated in our samples. The pathways identified were then contrasted with our full dataset, and we derived counts for the genes that were either over- or under-expressed for each pathway of interest in each one of our conditions. To account for biases in annotation, as well as for the difference in gene counts for each pathway, we chose to express our data as a ratio of genes that are differentially expressed by the total amount of genes associated with a given pathway. The raw reads as well as the resulting transcript count matrix were deposited at GEO, under accession number GSE252562.

Statistical analyses and data plotting

All statistical analyses were performed using R(*56*). For statistical analyses among survival curves, we considered each curve as a replicate rather than doing multiple comparisons for each timepoint. To accomplish this, we first estimated the area under the curve for each replicate using the “bayestestR” package(*59*). The AUC values were then compared with One-Way or Two-Way ANOVAs, depending on the experimental design. Datasets that were deemed significant by the ANOVA were then compared with two-sided pairwise t-tests (“stats” package(*56*)), with Bonferroni-Holm correction. Robust linear regression lines were fit with the “ggpmisc” package, using the stat_poly_line function(*60*). All data shown were plotted with R, using the “ggplot2”(*61*), “ggthemes”(*62*), and “ggbeeswarm”(*63*) packages. Final figures were assembled with the Adobe Illustrator software. For all figures in which “survival” is plotted, the average and original data points are slightly offset to facilitate visualization, but were all collected at the timepoints indicated.

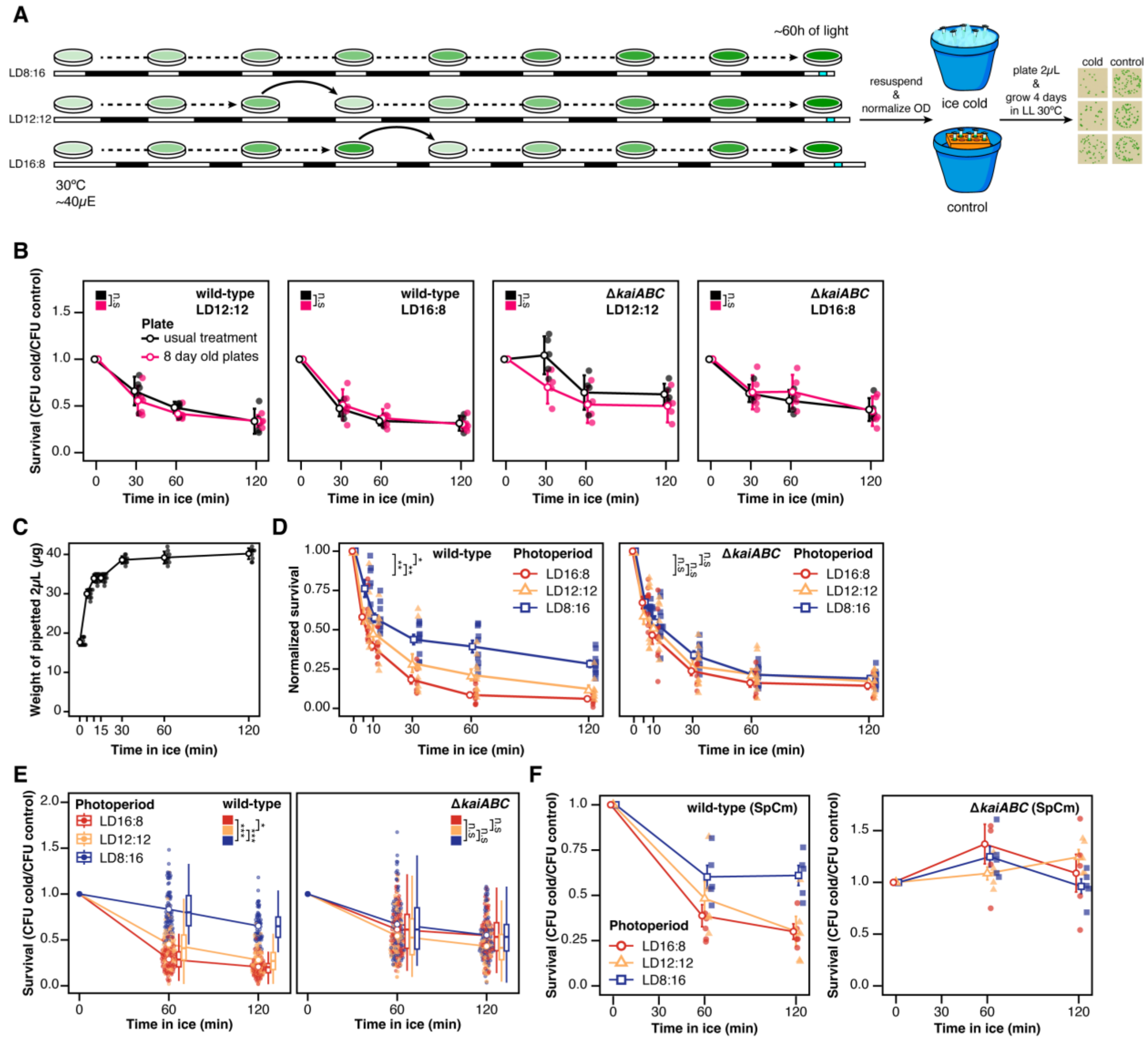


**Fig. S1. Methodology for the cold survival assay.**

(**A**) Plates inoculated with cyanobacteria were exposed to different photoperiods (here, LD8:16, LD12:12 and LD16:8) for a total of 8 days at 30ºC and ~40 μE $m^{-2}$ $s^{-1}$ (see Material and Methods for further information about growth conditions). (**B**) To test whether our replating protocol for LD16:8 and LD12:12 had any effect in our results, we ran our assay in cells that were given the same treatment as shown in (A), and cells that were transferred to plates which were put in the experimental chamber at the same time as the LD8:16 samples, but were only inoculated at the same time as the replatings in our usual protocol ("8 day old plates;" *n* = 5 for all except Δ*kaiABC* LD12:12 "usual treatment" and WT LD12:12 "8 day old plates" 120min, in which *n* = 4). No difference among treatments was observed for either photoperiod in wild-type and Δ*kaiABC* (pairwise t-tests with Bonferroni correction). (**C**) Due to the difference in temperature of the experimental room (30ºC) and the bacterial suspensions, the actual weight of the 2μL pipetted throughout the survival assay varied(*49*). Measurements of weight at each timepoint were used to normalize the survival curves in panel d (*n* = 11 for 0min, 10min and 15min; *n* = 9 for 5min, *n* = 10 for 30min, 60min and 120min). (**D**) Survival curves for wild-type (*n* = 10 for all except LD8:16 5min, in which *n* = 8) and Δ*kaiABC***,** normalized according to c (*n* = 9 for LD16:8; *n* = 10 for LD12:12 except for 10min in which *n* = 9; *n* = 10 for LD8:16 except 5min and 120min in which *n* = 9). The data plotted here are the same as in Fig. 1B. (**E**) Combination of ALL the experiments shown throughout this manuscript in which wild-type and Δ*kaiABC* cells were tested after 8 cycles of LD16:8 (*n* = 79 and 74 for wild-type, 60 and 120min; *n* = 77 and 72 for Δ*kaiABC*, 60 and 120min), LD12:12 (*n* = 74 and 65 for both wild-type and Δ*kaiABC*, 60 and 120min) or LD8:16 (*n* = 79 and 74 for wild-type, 60 and 120min; *n* = 79 and 73 for Δ*kaiABC*, 60 and 120min). Open dots within the cloud of dots show the average, and the bars show the standard error of the mean. Boxplots show the distribution of the data. (**F**) To confirm that the differing antibiotic resistances of the wild-type and the Δ*kaiABC* strains used throughout the experiments were not responsible the differing responses to photoperiod, we repeated the photoperiodic assay in wild-type and Δ*kaiABC* that had been transformed to be resistant to both spectynomicin (Sp) and chloramphenicol (Cm); *n* = 5 except for Δ*kaiABC* LD16:8 at 60min, in which *n* = 4. Asterisks refer to significance (* = <0.05, ** = < 0.01) in a pairwise t-test with Bonferroni correction. For all survival curves, the open symbols denote the average and the error bars denote the standard error of the mean.

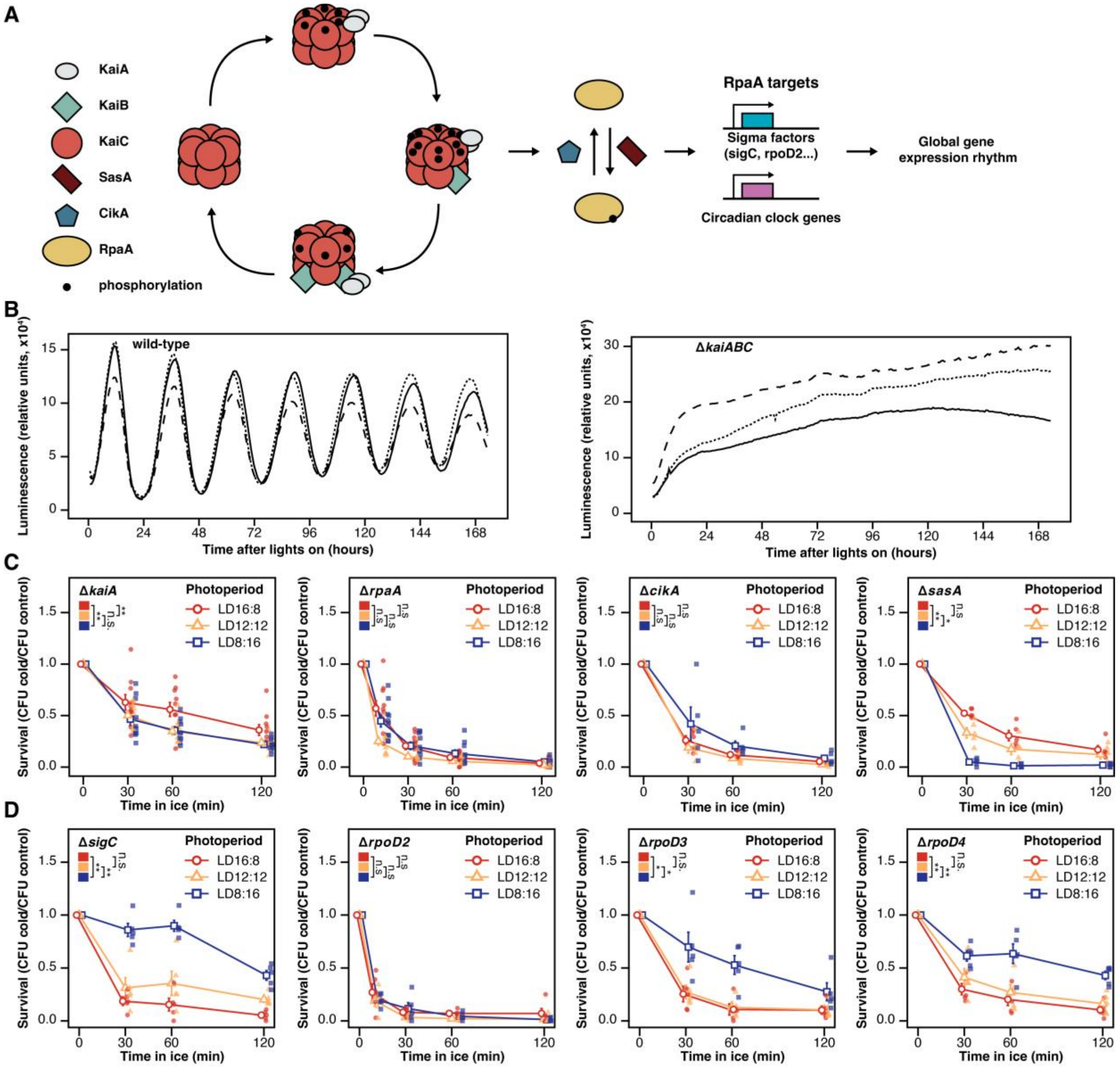


**Fig. S2. Disruption of the circadian clock leads to loss of photoperiodic response.**

(**A**) Diagram of the circadian clockwork in cyanobacteria, showing the core proteins that compose the clock mechanism (left) as well as its downstream targets (right). (**B**) Rhythms of gene expression for wild-type and Δ*kaiABC* colonies assessed as rhythms of luminescence emitted by the P*kaiBC::luxAB* reporter**(*6*)**. 3 representative traces are shown, selected from at least 15 simultaneously measured replicates. (**C**) Photoperiodic assay (performed as described in Fig. S1) for knock-out mutants of *kaiA* (*n* = 10), *rpaA* (*n* = 10), *cikA* (*n* = 5) and *sasA* (*n* = 5). (**D**) Same as (C), but instead for knock-outs of the sigma factors *sigC* (*n* = 5), *rpoD2* (*n* = 5) (direct targets of RpaA**(*21*)**), *rpoD3* (*n* = 5) and *rpoD4* (*n* = 5) (indirect targets of RpaA). Asterisks refer to significance (* = <0.05, ** = < 0.01) in pairwise t-tests with Bonferroni correction. For all survival curves, the open symbols denote the average and the error bars denote the standard error of the mean.

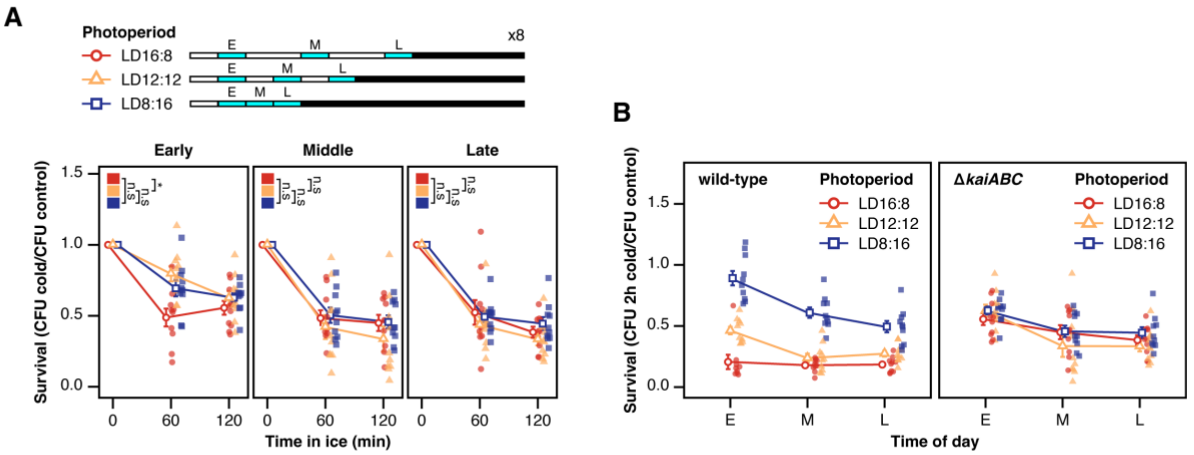


**Fig. S3. The Δ*kaiABC* clock-knockout mutant does not show a photoperiodic response regardless of the time of day analyzed.**

(**A**) Similarly to Fig. 1C, Δ*kaiABC* cells were measured for cold sensitivity after exposure to either short, equinox or long days, at the beginning (2h after dawn), middle (midpoint) and end (2h before dusk) of the day ($n$ = 10 for all). Differences were tested for significance with a pairwise t-test with Bonferroni correction. (**B**) Survival after cold shocks (shown here after 2h) is higher during the beginning of the day and lower at the end of the day for wild-type cells exposed to short-days. On the other hand, Δ*kaiABC* cells coming from different photoperiods show much smaller/no dependence on time of day of the assay. For all survival curves, the open symbols denote the average and the error bars denote the standard error of the mean

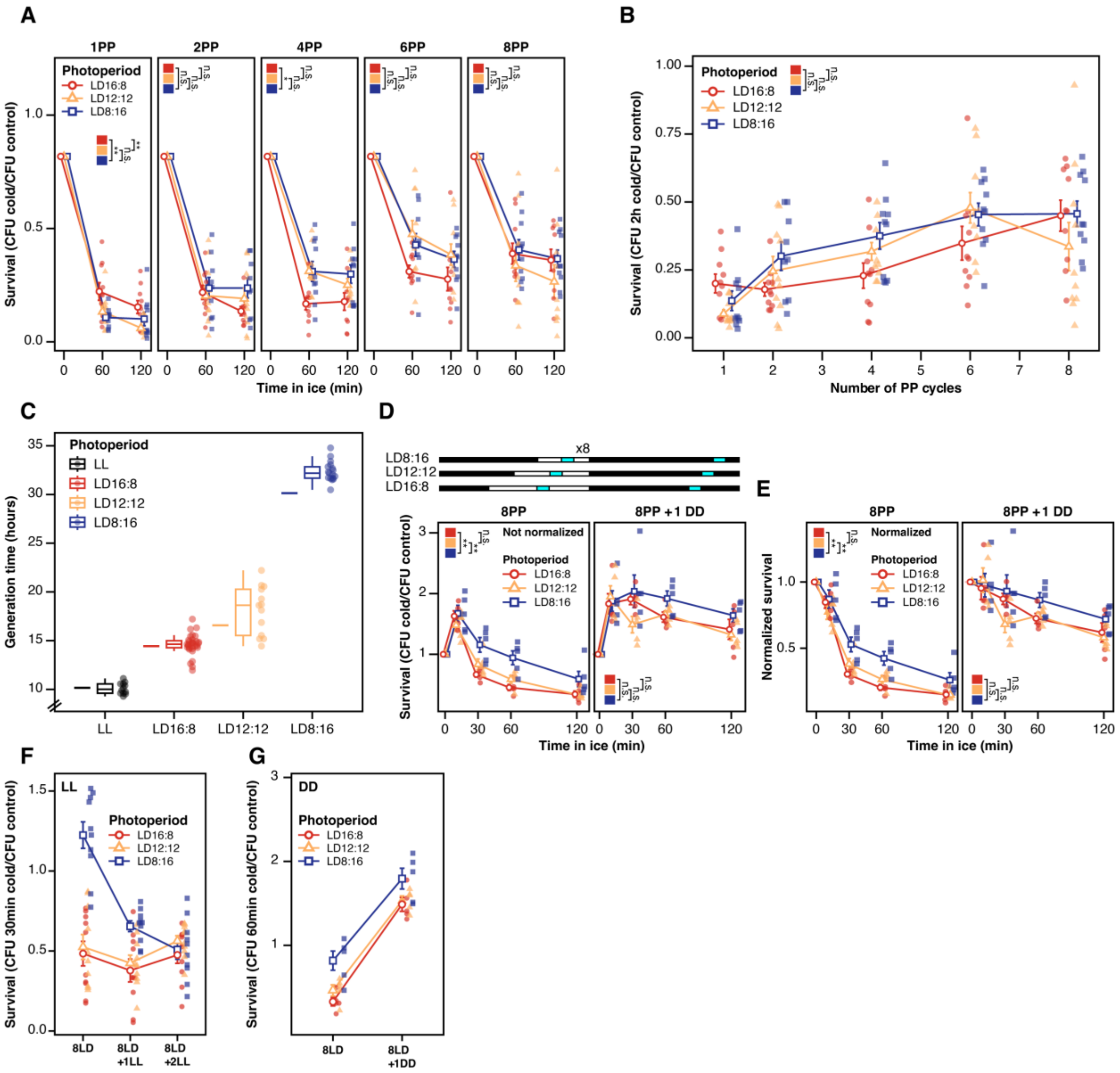

**Fig. S4. Further tests of history dependence.**

(**A**) Survival curves similar to Fig. 2A, but showing results for Δ*kaiABC* cells. (**B**) Survival after 2h of exposure to cold throughout the different number of PP cycles (*n* = 10 for all except LD8:16 at 2 PP, for which *n* = 9). (**C**) Generation time for each of the photoperiods analyzed, as well as LL (from left to right, *n* = 12, 31, 13, 17). Bars to the left of the boxplots show the median generation time for wild-type cells exposed to the same photoperiods. (**D**) Diagram showing the protocol for the constant darkness experiments, with blue bars indicating the time in which the cold survival assay was performed, and survival curves showing the results of the experiment (*n* = 5). (**E**) The same data as in (D), but normalized using the conversion shown in fig. S1C. (**F**) Comparison of survival after 30min of cold exposure for the constant light experiment. (**G**) Similar to (F), but showing survival after 60min of cold for the constant darkness experiment. For all survival curves, the open symbols denote the average and the error bars denote the standard error of the mean. Significance was determined through pairwise t-tests with Bonferroni correction.

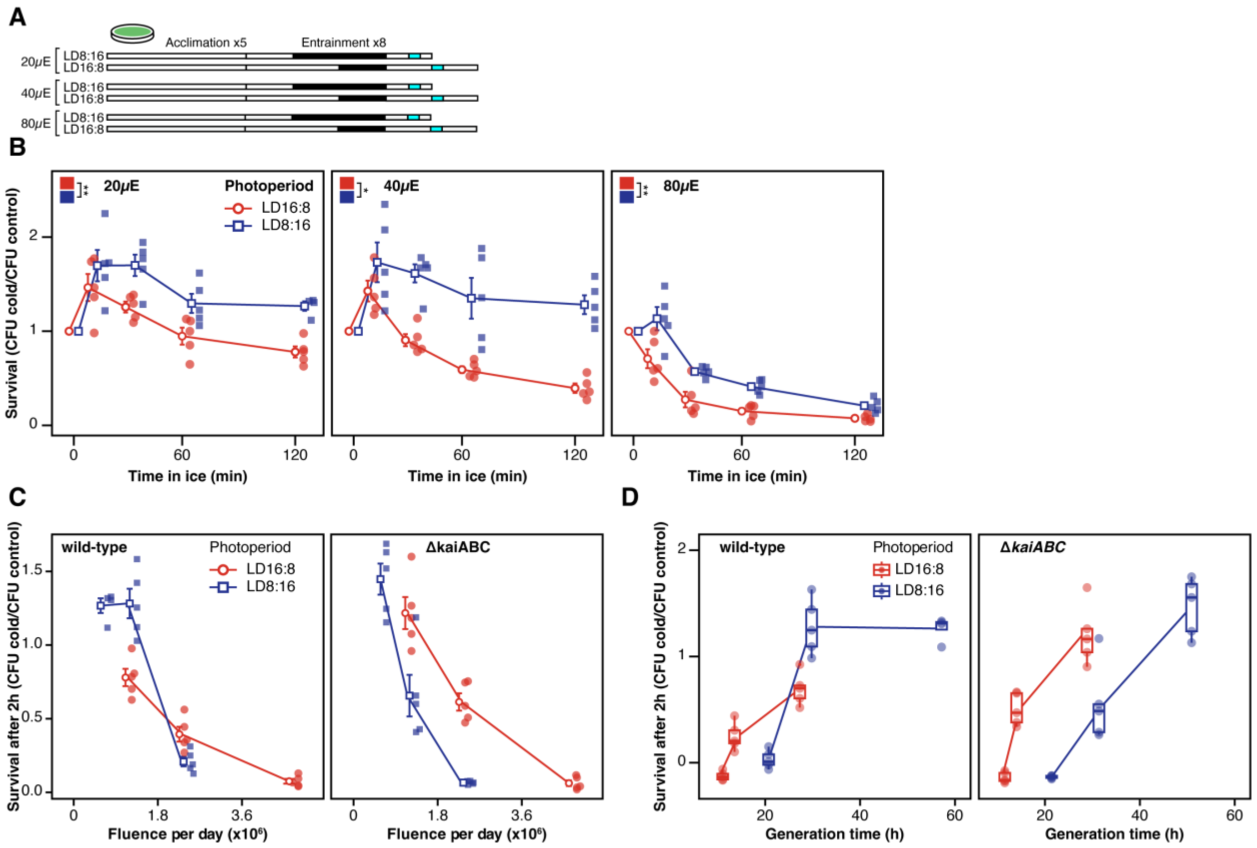


**Fig. S5. Light intensity influences the amplitude of the cold photoperiodic response.**

(**A**) Diagram showing the protocol for the different light intensities and photoperiods tested. Blue bars indicate the time in which the cold survival assay was performed (**B**) Survival curves for wild-type cells exposed to different fluences ("µE" = µE $m^{-2}$ $s^{-1}$); $n$ = 5 for all except 20µE LD8:16 at 120min, in which $n$ = 4). (**C**) The data in panel b were replotted to show how survival after 2h of cold exposure varied as a function of the total fluence in a day (calculated by multiplying the light intensity for the amount of second of light in each day) ($n$ = 5 for all except LD8:16 at the lowest fluence, in which $n$ = 4). (**D**) The data in panel (B) and (C) were combined with growth data in fig. S18B to show how survival after 2h of cold exposure varied as a function of generation time ($n$ = 5 except for wild-type at 60h, in which $n$ = 4). For panels (B-D), the open symbols denote the mean +/- SEM, blue symbols are data from LD8:16 and red symbols are data from LD16:8.

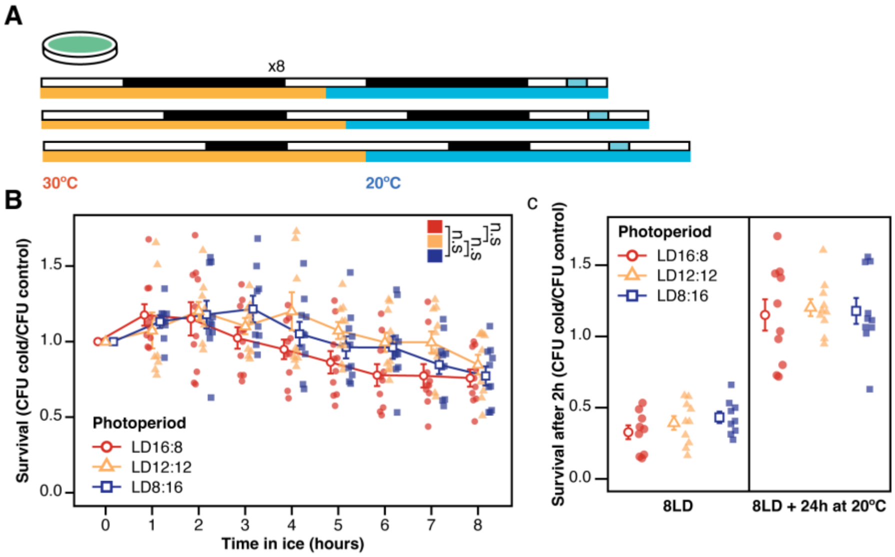


**Fig. S6. Δ*kaiABC* shows no difference in survival across different photoperiods with or without prior 20ºC exposure.**

(**A**) Diagram of the experiment testing the interaction of photoperiod and intermediate cold temperatures on survival. (**B**) Survival curve for Δ*kaiABC* cells (*n* = 10 for all except LD8:16 at 1h and 3h, in which *n* = 9, and LD12:12 1h, in which *n* = 8). (**C**) Comparison between the survival of Δ*kaiABC* cells after 2h of exposure to cold, with and without prior treatment of 24h at 20ºC.

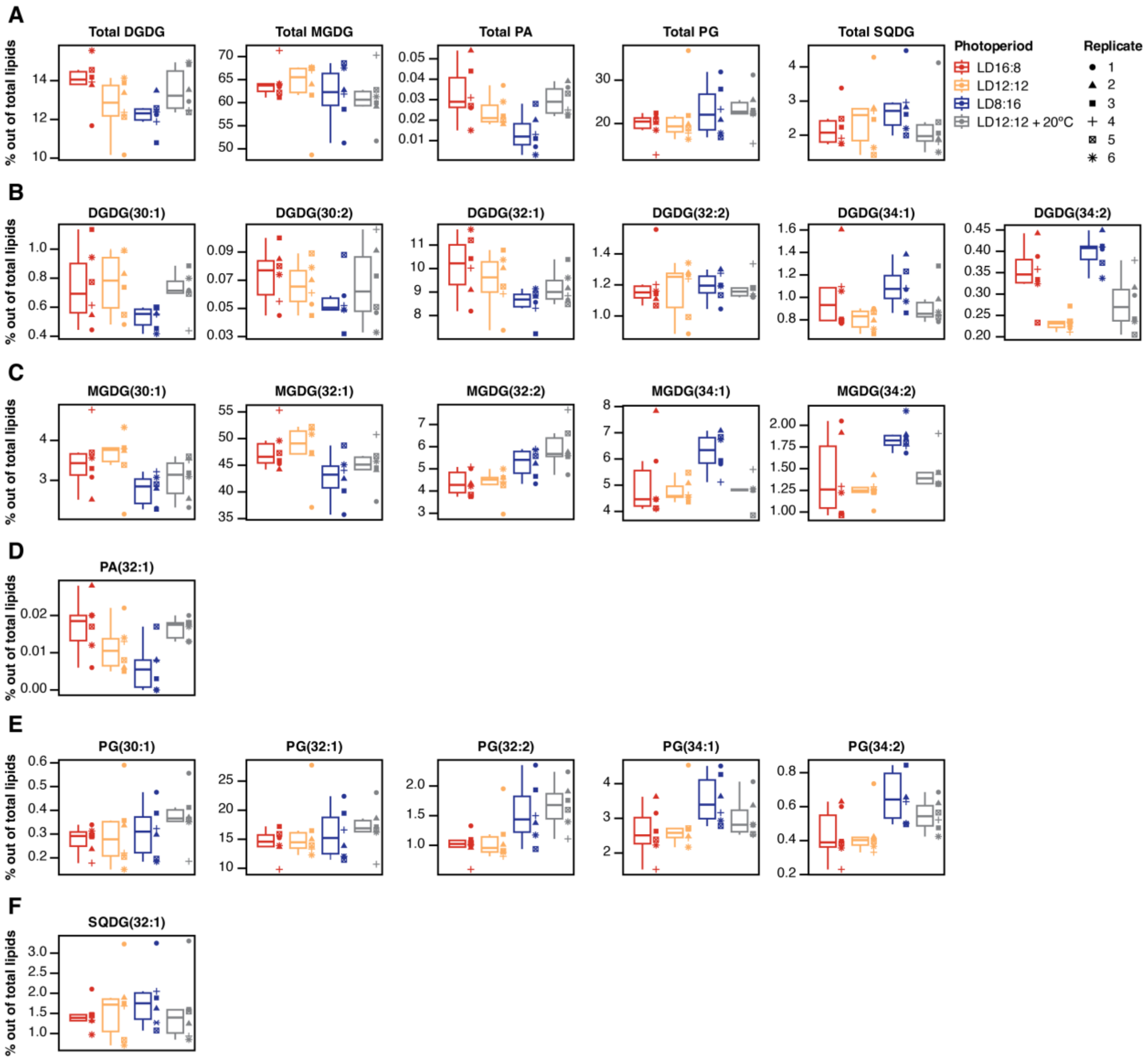


**Fig. S7. Amount of each lipid species and each lipid class, expressed as a percentage of the total lipids for wild-type samples.**

(**A**) Percentages of each lipid class. (**B**) Percentages of identified DGDG species. (**C**) Percentages of identified MGDG species. (**D**) Percentages of identified PA species. (**E**) Percentages of identified PG species. (**F**) Percentages of identified SQDG species. For all samples, *n* = 6. Abbreviations: MGDG = monogalactosyldiacylglycerol; DGDG = digalactosyldiacylglycerol; PG = phosphatidylglycerol; SQDG = sulfoquinovosyl diacylglycerol; PA = phosphatidic acid.

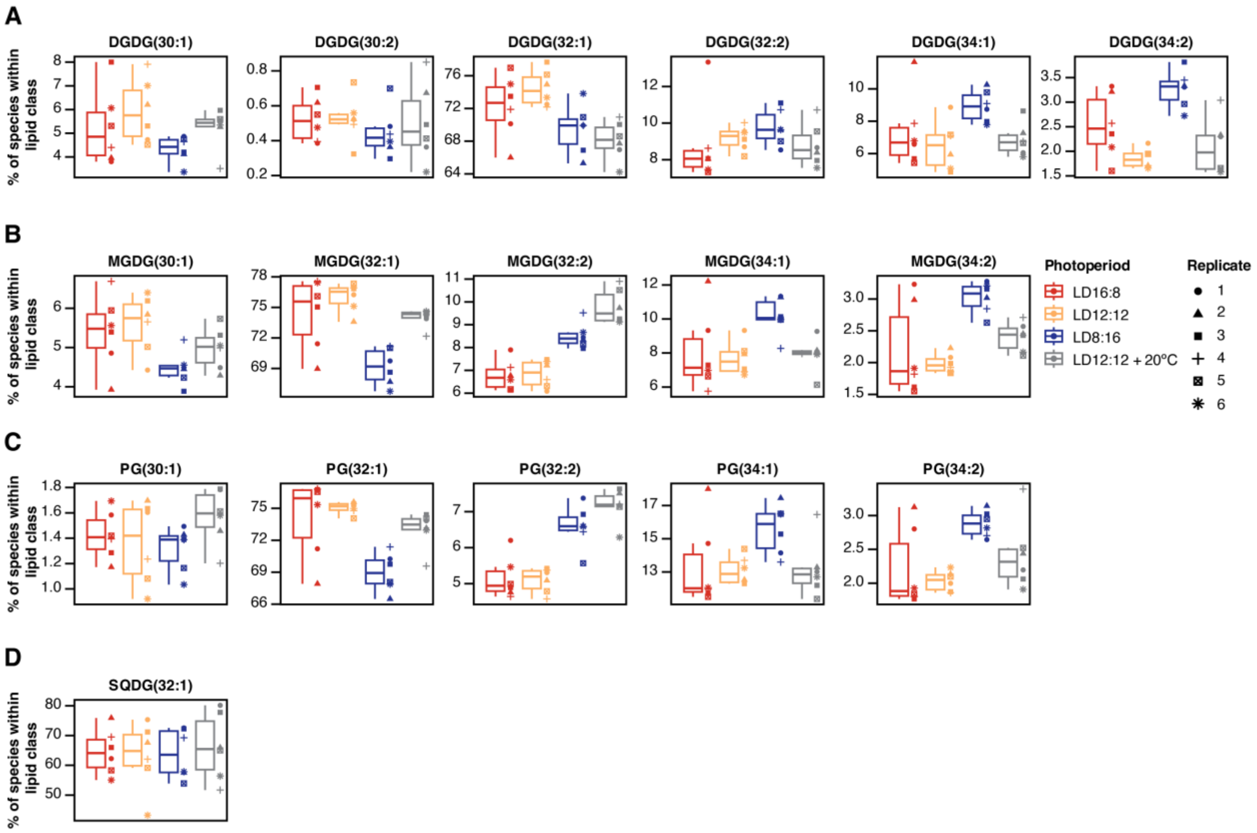


**Fig. S8. Amount of each lipid species expressed as a percentage of the total of each lipid class, for wild-type samples.**

(**A**) Percentages of identified DGDG species. (**B**) Percentages of identified MGDG species. (**C**) Percentages of identified PG species. (**D**) Percentages of identified SQDG species. For all samples, $n = 6$. Abbreviations of lipid species as in fig. S7.

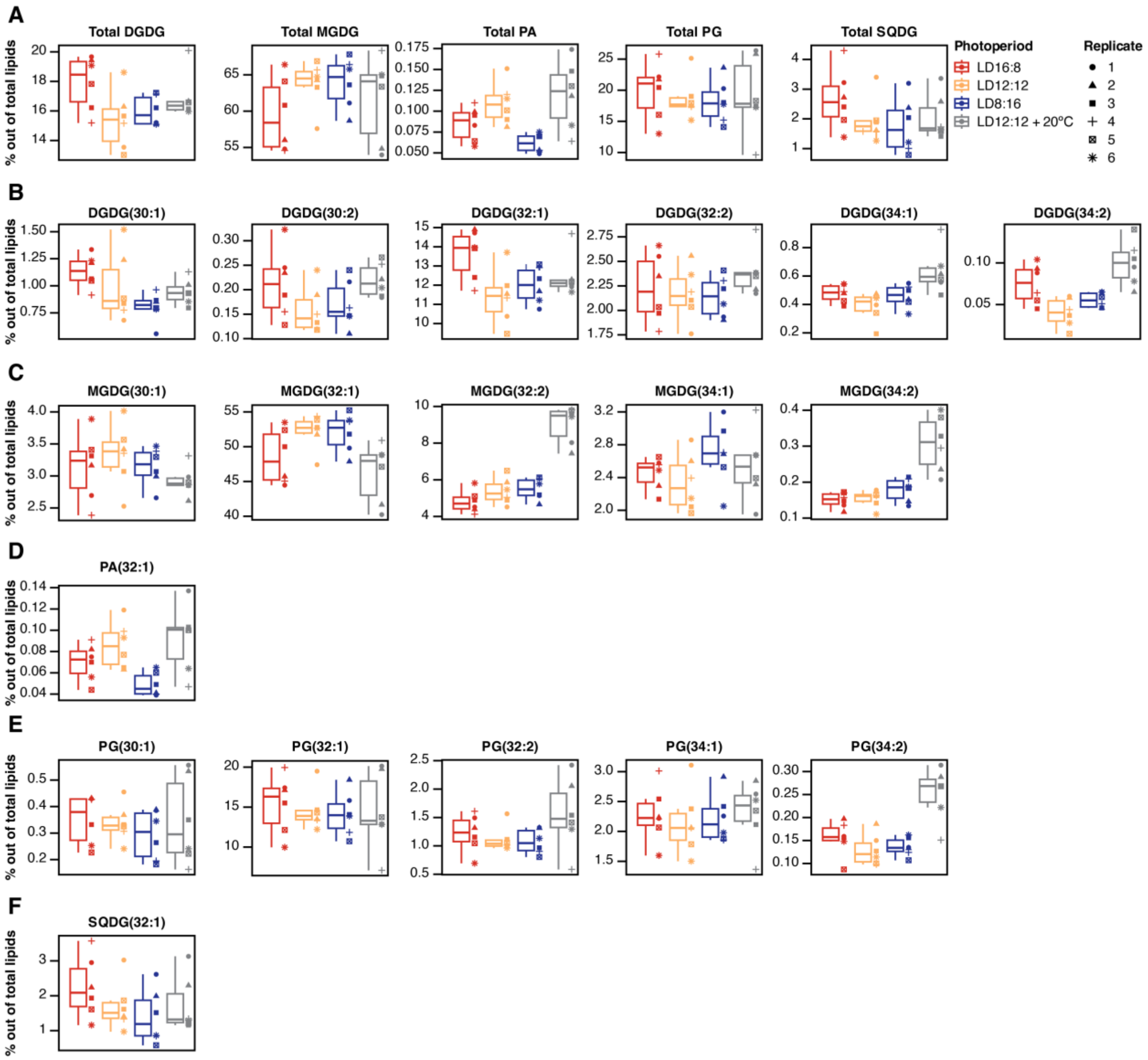


**Fig. S9. Amount of each lipid species and each lipid class, expressed as a percentage of the total lipids for Δ*kaiABC* samples.**

(**A**) Percentages of each lipid class. (**B**) Percentages of identified DGDG species. (**C**) Percentages of identified MGDG species. (**D**) Percentages of identified PA species. (**E**) Percentages of identified PG species. (**F**) Percentages of identified SQDG species. For all samples, *n* = 6. Abbreviations of lipid species as in fig. S7.

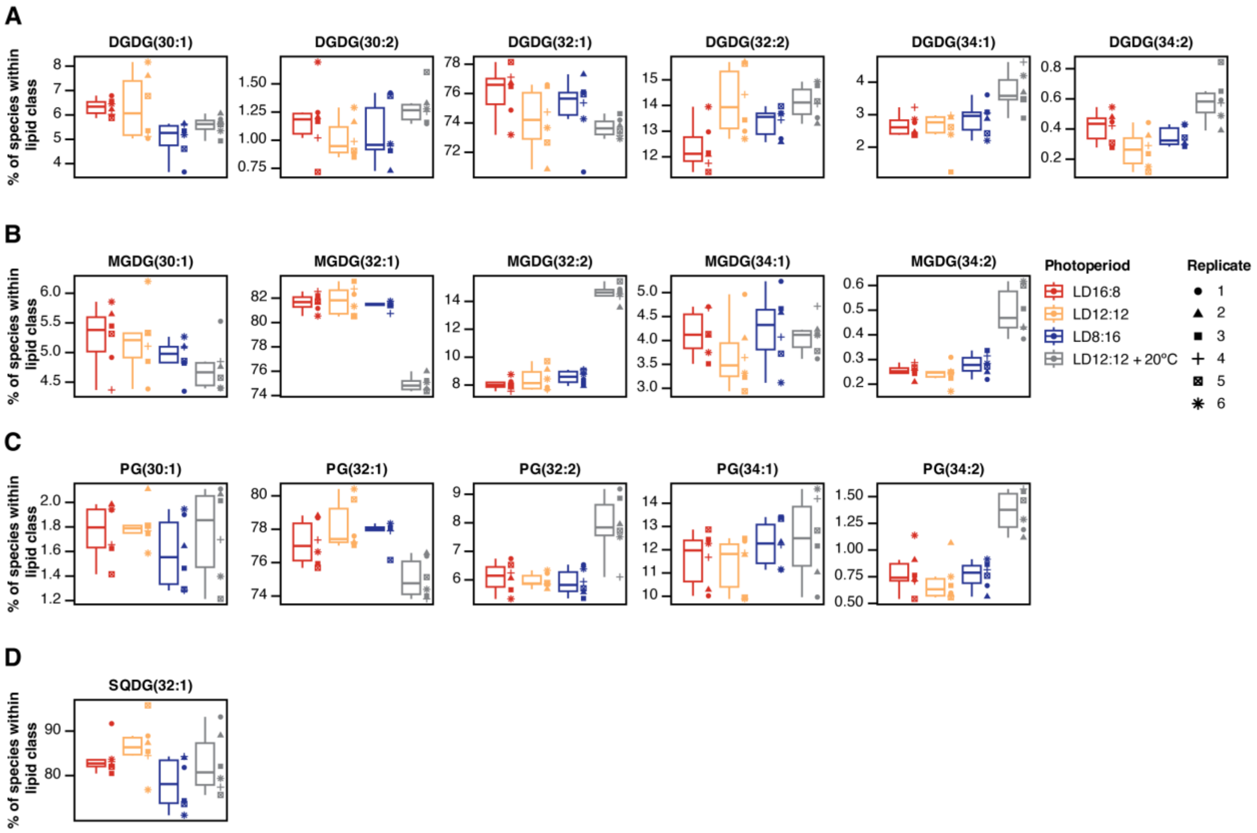


**Fig. S10. Amount of each lipid species expressed as a percentage of the total of each lipid class, for Δ*kaiABC* samples.**

(**A**) Percentages of identified DGDG species. (**B**) Percentages of identified MGDG species. (**C**) Percentages of identified PG species. (**D**) Percentages of identified SQDG species. For all samples, $n$ = 6. Abbreviations of lipid species as in fig. S7.

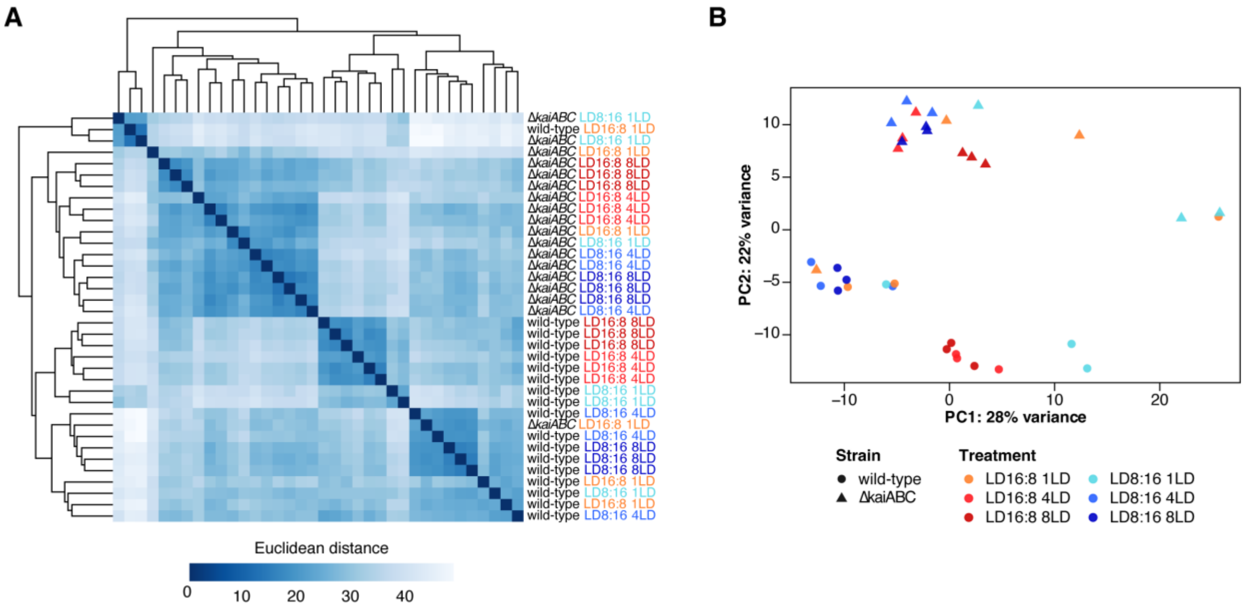


**Fig. S11. Summary of the RNA-seq samples.**

(**A**) Heatmap showing the hierarchical clustering of the RNA-seq samples, estimated by Euclidean distances. Red labels refer to long-day samples, and blue labels to short-day samples. (**B**) PCA of the same samples. The colors match those of panel (A).

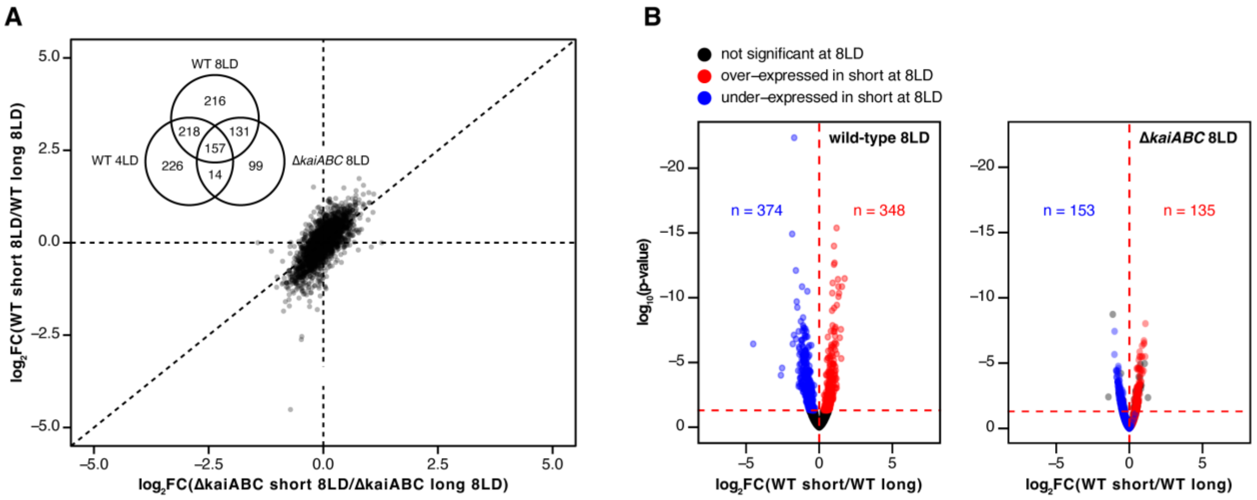


**Fig. S12. Overlap between differential expression in wild-type and Δ*kaiABC* cells exposed to 8 PP cycles.**

(**A**) Correlation plot showing the log2(fold change) in expression of short-day cells divided by long-day cells, comparing wild-type cells exposed to 8 PP to Δ*kaiABC* cells exposed to 8 PP. The Venn diagram inside the correlation plot shows the number of differentially expressed genes shared between wild-type cells at 8 PP and 4 PP, as well as Δ*kaiABC* cells at 8 PP. Numbers in parentheses indicate the total number of genes differentially expressed for that group. They do not include genes that are differentially expressed in opposite ways. The vertical and horizontal dashed lines indicate the y and x zero intercepts, and the diagonal line shows y=x. (**B**) Volcano plots showing the differentially expressed genes for wild-type at 8 PP (left) and Δ*kaiABC* at 8 PP (right), colored by the difference in expression for wild-type at 8 PP. The numbers in blue on the left plot refer to the total number of genes under-expressed in short-days for wild-type after 8 PP, while those in red show the genes over-expressed. On the plot to the right, the numbers refer to genes under or over-expressed in Δ*kaiABC* after 8 PP that are also under or over-expressed in wild-type after 8 PP.

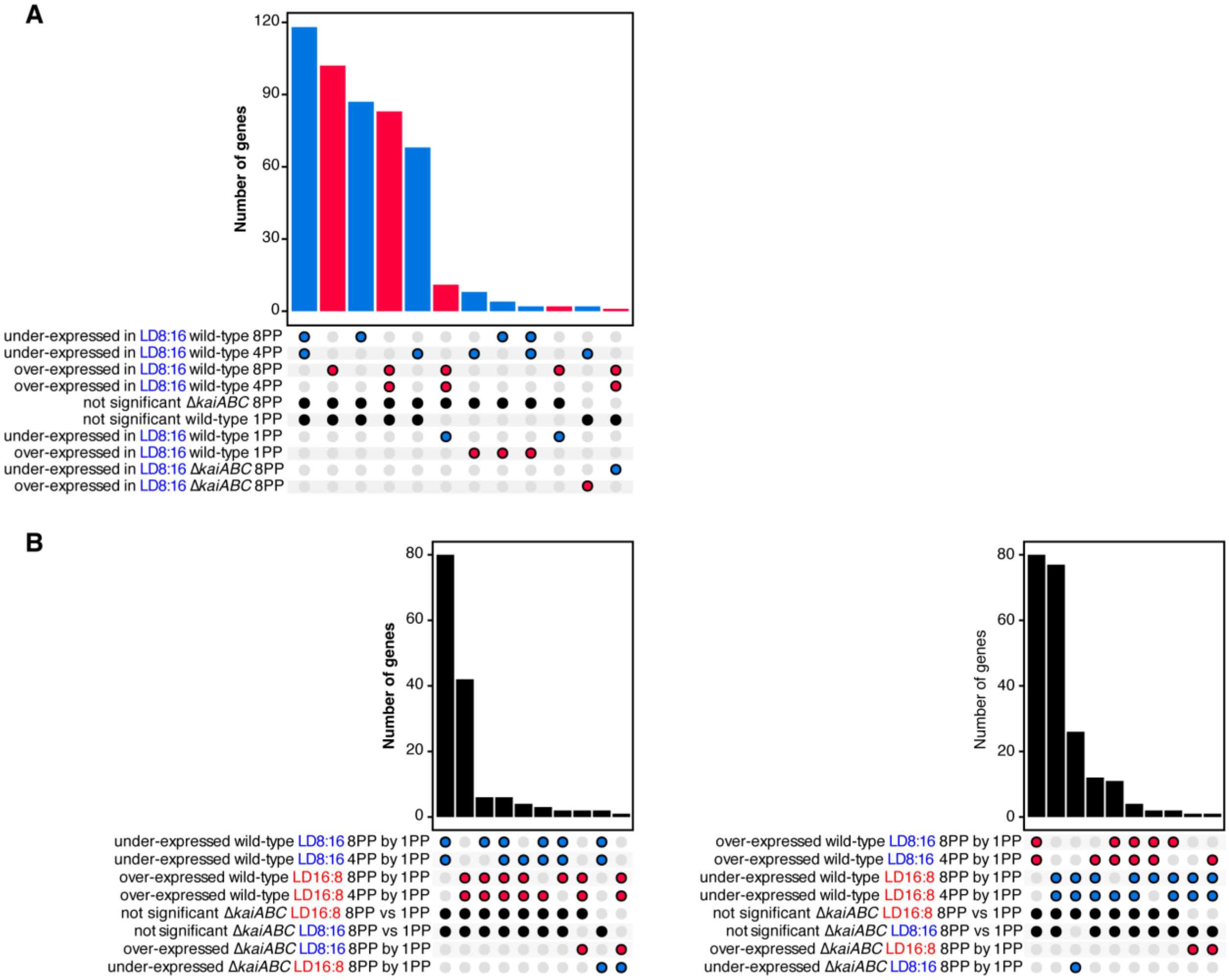


**Fig. S13. UpSet plots showing our genes of interest.**

(**A**) Genes of interest obtained comparing LD8:16 samples to LD16:8 samples within the same number of PP cycles ($n$ = 488). Blue bars indicate genes that are under-expressed in LD8:16 in either 8 PP or 4 PP, and red bars indicate genes those that are over-expressed. (**B**) Genes of interest obtained comparing samples of the same photoperiod across different numbers of PP ($n$ = 364; 145 genes are common to both (A) and (B). For all UpSet plots, blue dots denote under-expression at the indicated group, and red dots denote over-expression.

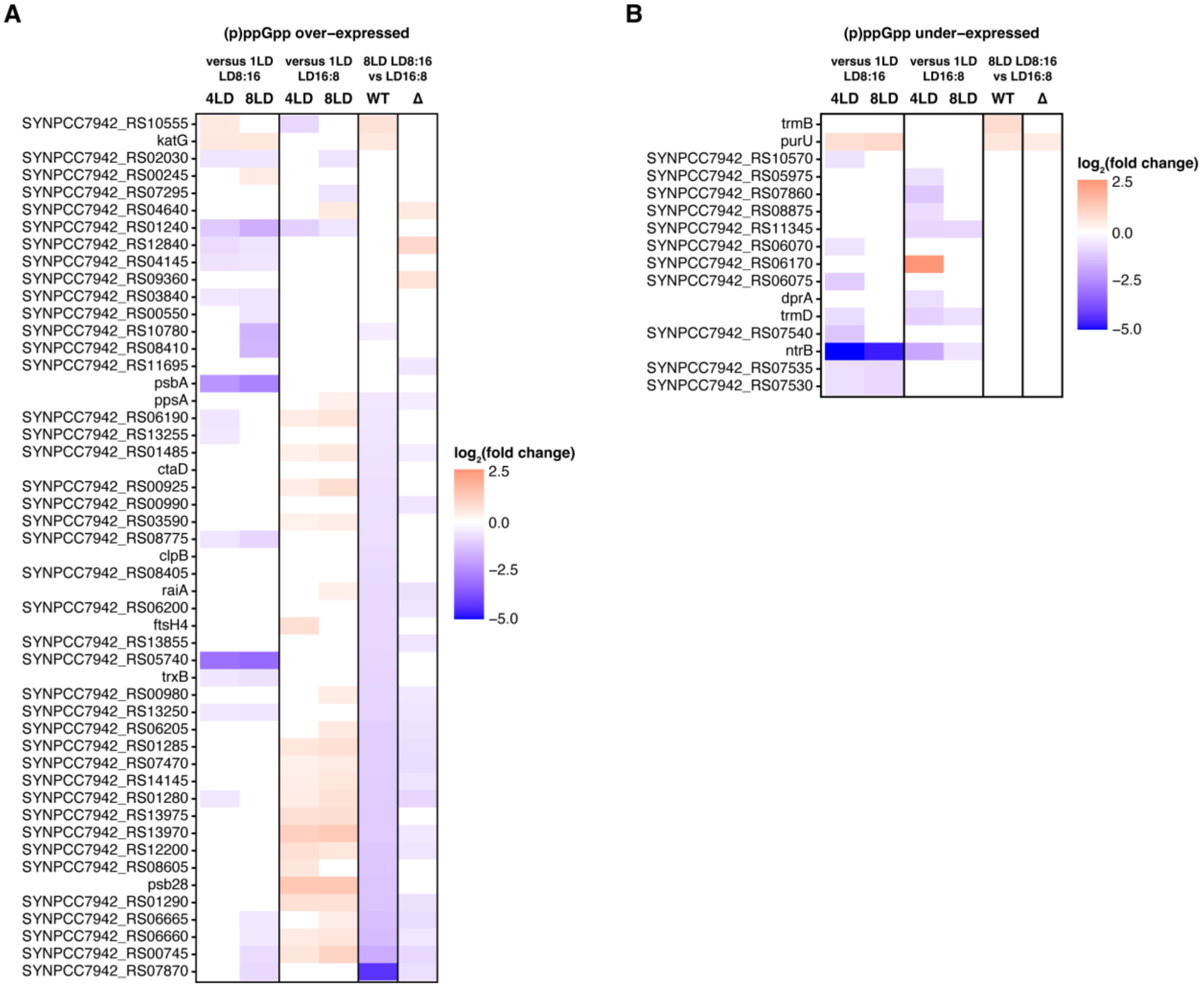


**Fig. S14. Genes related to the stringent response show differences in expression between short and long days in wild-type.**

(**A**) Heatmap showing genes that were shown to be over-expressed by the stringent response ((p)ppGpp over-expressed)***(30)***. The genes are ordered by their log2(fold change) in wild-type, for comparisons between 8 PP LD8:16 and 8 PP LD16:8. Similar to Fig. 4D, the first four columns show the $\log_2$(fold-change) in expression in wild-type cells within each photoperiod of either 4 PP or 8 PP against 1 PP. The last two columns show the $\log_2$(fold-change) for 8 PP 8:16 vs 8 PP 16:8, for wild-type (fifth column, WT) and Δ*kaiABC* (sixth column, Δ). (**B**) Similar to panel (A), but showing genes that were shown to be under-expressed by the stringent response ((p)ppGpp under-expressed).

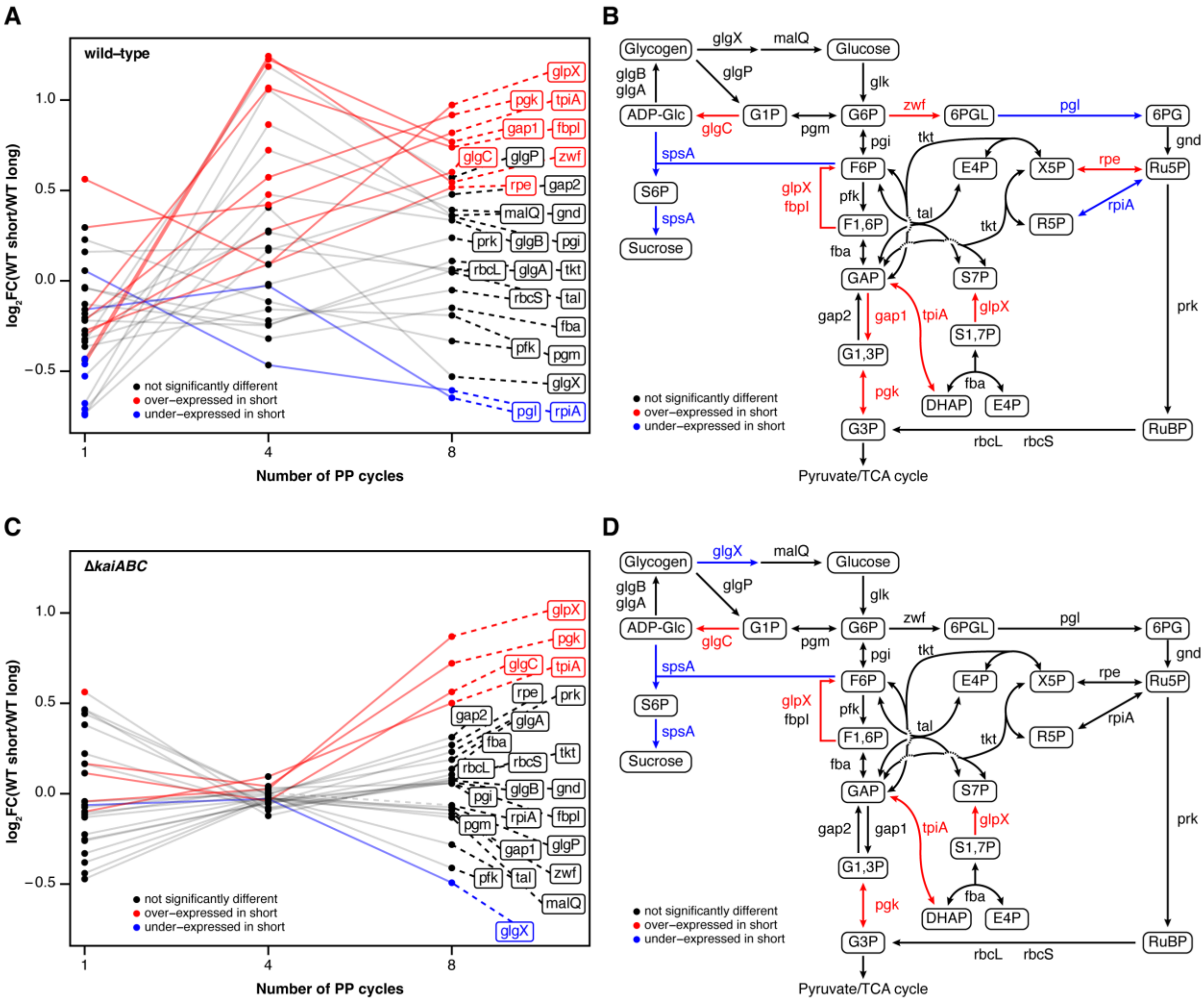


**Fig. S15. Effects of long and short days on genes related to important metabolic pathways.** (**A**) Plot showing the $\log_2$ fold-change in expression between short-day wild-type and long-day wild-type across different numbers of PPs. The genes shown are involved in glycogen and sucrose metabolism, glycolysis, the Calvin Cycle and the oxidative pentose phosphate pathway. Dots are colored based on their differential expression at the PP indicated, while lines are trajectories that are colored based on the differential expression for each gene at 8 PP. Dashed lines connect the labels of each gene with the data points at 8 PP. Gene labels are also colored based on their differential expression at 8 PP. (**B**) Diagram showing the glycogen metabolism pathway, glycolysis, Calvin cycle and the oxidative pentose phosphate pathway, including the genes related to each enzymatic step. Genes are colored based on their differential expression for wild-type at 8 PP, with genes labeled in blue being significantly under-expressed in short, genes in red being significantly over-expressed, and those in black being not differentially expressed. (**C**) Similar to a, but showing the patterns for the Δ*kaiABC* strain (i.e., $\log_2$ fold-change in expression between short-day Δ*kaiABC* and long-day Δ*kaiABC*. (**D**) Similar to b, but showing the patterns for the Δ*kaiABC*. ADP-Glc = ADP-glucose; DHAP = dihydroxyacetone phosphate; E4P = erythrose-4-phosphate; F1,6P = fructose-1,6-bisphosphate; F6P = fructose-6-phosphate; GAP = glyceraldehyde-3-phosphate; G1P = glucose-1-phosphate; G1,3P = 1,3-

bisphosphoglycerate; G3P = 3-phosphoglycerate; G6P = glucose-6-phosphate; RuBP = ribulose-1,5-bisphosphate; Ru5P = ribulose-5-phosphate; R5P = ribose-5-phosphate; S7P = sedoheptulose-7-phosphate; S1,7P = sedoheptulose-1,7-bisphosphate; X5P = xylose-5-phosphate; 6PG = 6-phosphogluconate; 6PGL = 6-phosphogluconolactone.

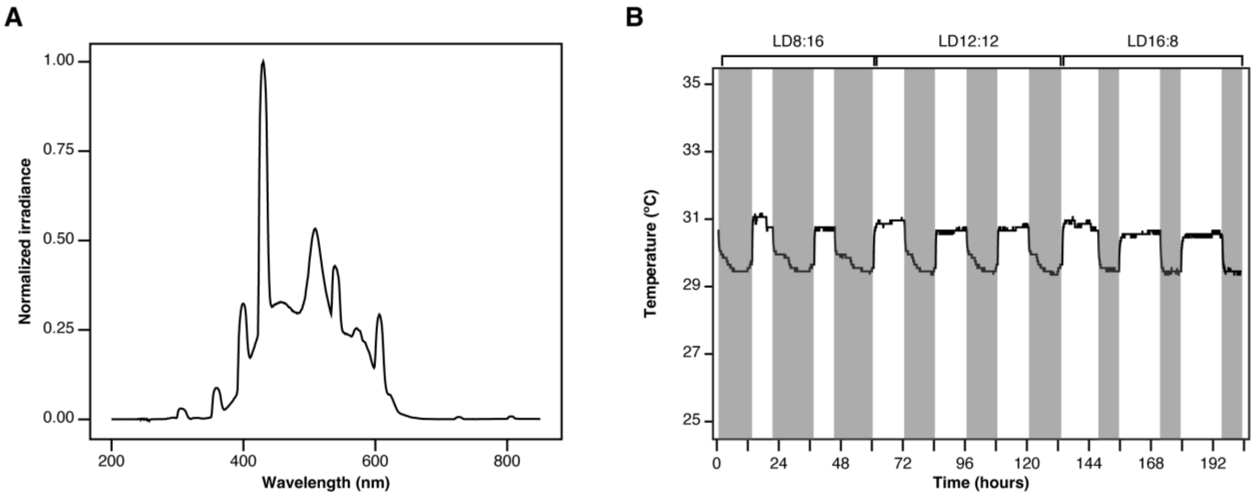


**Fig. S16. Light and temperature conditions used throughout most of the experiments described.**

(**A**) Light spectrum of the fluorescent lamp used. (**B**) Temperature measurements for the three main photoperiods used in the paper. Grey intervals refer to the dark phase, while white intervals refer to the light phase.

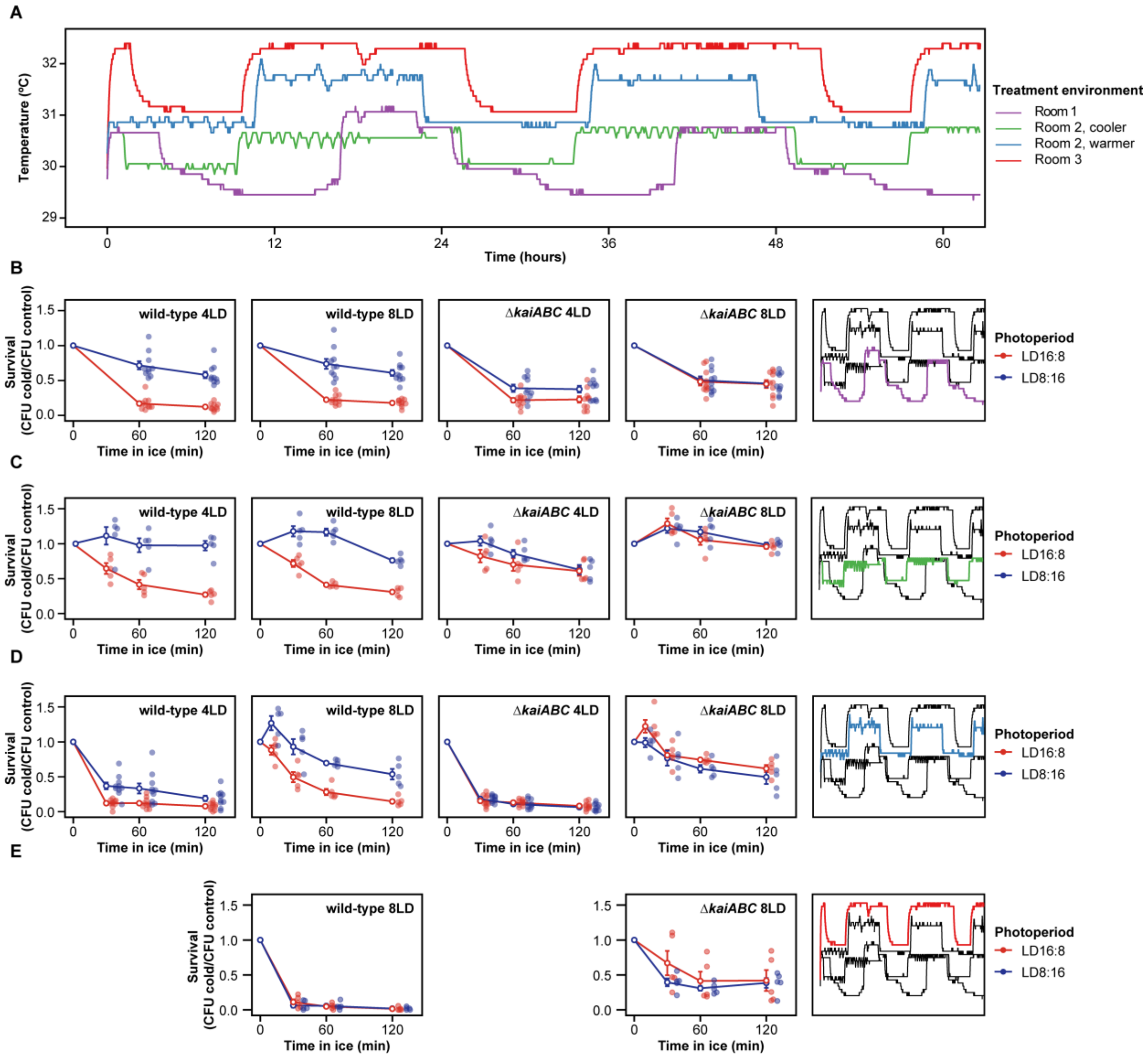


**Fig. S17. Temperature affects the amplitude of the photoperiodic response**

(**A**) Daily temperature variation in the three culture rooms and conditions tested. The results included in the rest of the paper come from "Room 1" and "Room 2, cooler" (**B**) Survival curves for wild-type and Δ*kaiABC* cells after 4LD and 8LD in "Room 1" (**C**) Same as (B), but for "Room 2, cooler" (**D**) Same as (B), but for "Room 2, warmer" (**E**) Same as (B), but for "Room 3" and only 8LD. The right-most plot of panels (B)-(E) shows a reproduction of the temperature profile in panel (A), highlighting the room/condition in which the assays were performed. For all plots, open symbols denote the average and the bars show the standard error of the mean. Actual data points are slightly offset to facilitate visualization.

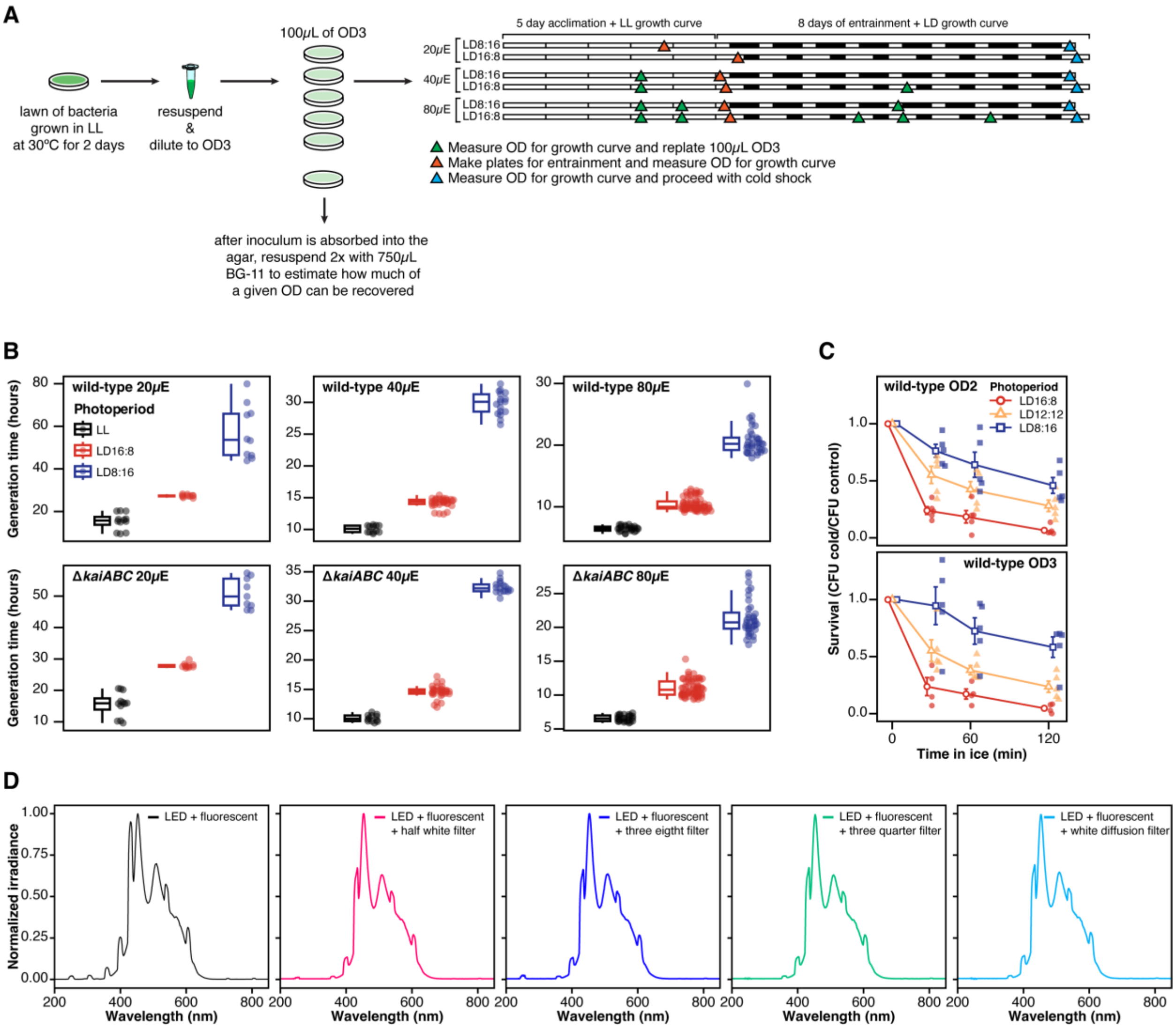


**Fig. S18. Protocol for light intensity experiments.**

(**A**) Diagram showing the preparation of the samples for the light intensity cold survival and growth curve experiments, which included a 5 day acclimation interval, followed by the usual 8 days of entrainment. (**B**) Generation times for wild-type (top) (from left to right, $n$ = 12, 10, 10, 12, 32, 17, 24, 68, 36) and Δ*kaiABC* (bottom) cells (from left to right, $n$ = 12, 10, 9, 12, 31, 17, 24, 66, 36) exposed to different photoperiods and light intensities. (**C**) Survival curves for cells that, unlike our usual streaking preparation, were inoculated on plates as 100µL of cell suspensions of either $OD_{750}$ = 2 ($n$ = 5) or $OD_{750}$ = 3 ($n$ = 5 except for LD16:8 in which $n$ = 4). All other methods were as shown in the diagram in fig. S1A. (**D**) Light spectrum of the lamps used in the light intensity experiments (fluorescent + LED), as well as the filters used to reduce the intensity to 80, 40 and 20µE $m^{-2}$ $s^{-1}$. For panel (C), the open symbols denote the average and the error bars denote the standard error of the mean.

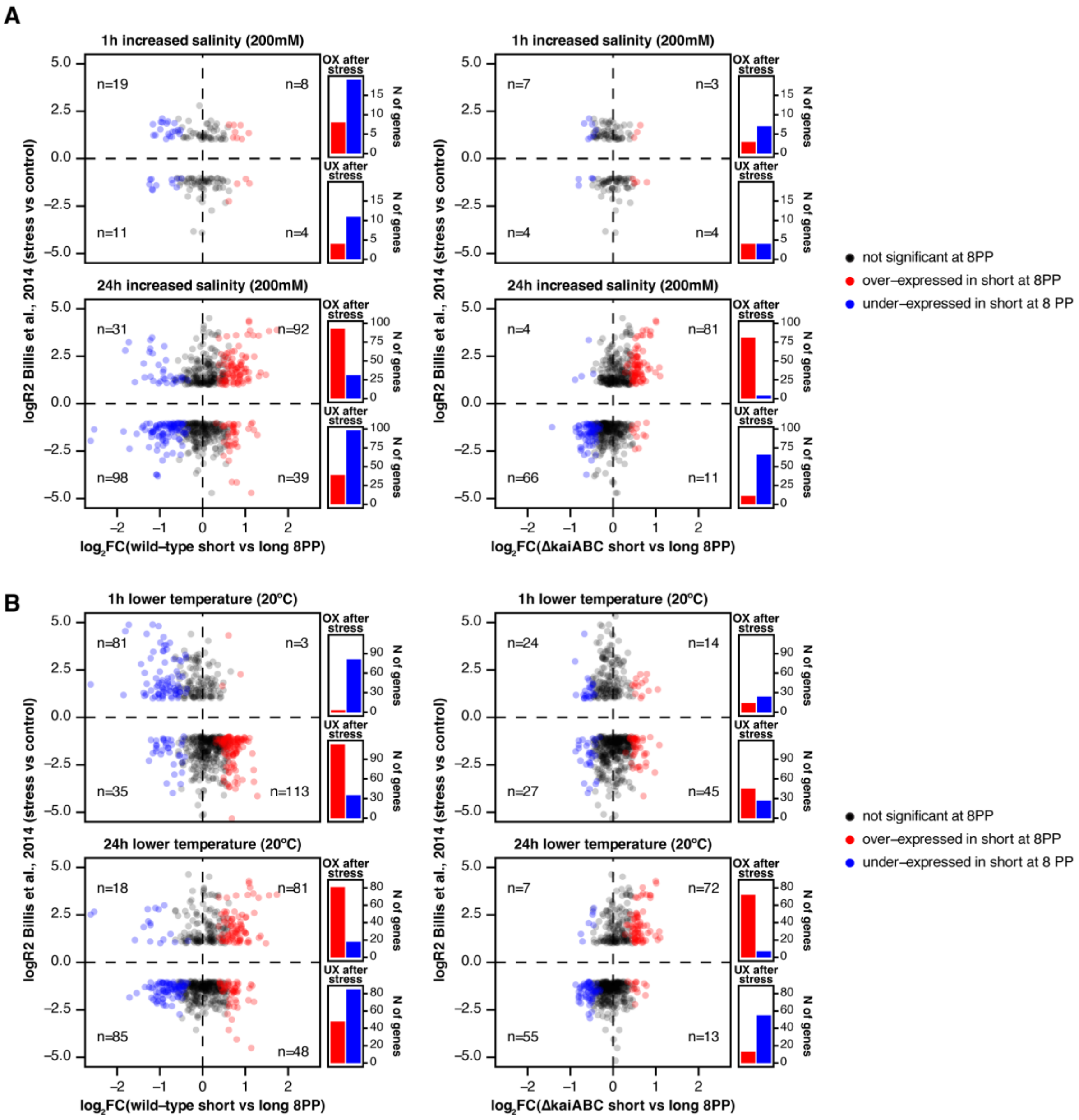


**Fig. S19. Correlation between differential gene expression after exposure to high salinity/low temperature stress and 8 PP of short days.**

(**A**) Differential expression of wild-type *S. elongatus* PCC 7942 after 1h and 24h of exposure to increased salinity (200mM of NaCl), independently produced(*40*), is contrasted against the differential expression between wild-type (right) or Δ*kaiABC* (left) after 8 PP of short vs long days. Only genes significantly different after increased salinity exposure are shown. Numbers refer to genes differentially expressed in short-days in each quadrant. (**B**) Similar to (A), but showing genes differentially expressed after 1h or 24h of transfer from 30ºC to 20ºC(*40*). Histograms show the number of genes that are over-expressed or under-expressed in LD8:16 at 8 PP, divided by whether they were over-expressed after stress (top) or under-expressed (bottom).
